# *In situ* cell-surface conformation of the TCR-CD3 signaling complex

**DOI:** 10.1101/2022.02.07.479368

**Authors:** Aswin Natarajan, Wenjuan Wang, Manuel Becerra Flores, Tianqi Li, Hye Won Shin, Saikiran Beesam, Timothy Cardozo, Michelle Krogsgaard

## Abstract

T cells play a vital role in adaptive immune responses to infections, inflammation and cancer and are dysregulated in autoimmunity. Antigen recognition by T cells – a key step in adaptive immune responses – is performed by the T cell receptor (TCR)-CD3 complex. The extracellular molecular organization of the individual CD3 subunits (CD3δε and CD3 γε) around the αβTCR is critical for T cell signaling. Here, we incorporated unnatural amino acid (UAA) photo-crosslinkers at specific mouse TCRα, TCRβ, CD3δ and CD3γ sites, based on previous mutagenesis, NMR spectroscopy and cryo-EM evidence, and crosslinking allowing us to identify nearby interacting CD3 or TCR subunits on the mammalian cell surface. Using this approach, we show that CD3γ and CD3ε, belonging to CD3γε heterodimer crosslinks to Cβ FG loop and Cβ G strand, respectively and CD3δ crosslinks to Cβ CC’ loop and Cα DE loop. Together with computational docking, we identify that in *in situ* cell-surface conformation, the CD3 subunits exists in CD3ε’-CD3γ-CD3ε-CD3δ arrangement around the αβ TCR. This unconventional technique, which uses the native mammalian cell surface microenvironment, includes the plasma membrane and excludes random, artificial crosslinks, captures a dynamic, biologically relevant, cell-surface conformation of the TCR-CD3 complex, which is compatible with the reported static cryo-EM structure’s overall CD3 subunits arrangement, but with key differences at the TCR-CD3 interface, which may be critical for experiments in T cell model systems.

## Introduction

T cell receptors (TCRs), expressed on T cells, recognize antigenic peptides presented by major histocompatibility complexes (MHC) expressed on antigen-presenting cells (APCs) and signal through associated CD3 subunits, resulting in T cell immune response initiation (Krogsgaard and Davis, 2005). The αβTCR is a heterodimeric molecule, with each subunit possessing a variable (Vα, Vβ) domain, which recognizes antigen through its complementarity-determining regions (CDRs), and a constant domain (Cα, Cβ), which facilitates interactions with CD3 subunits (Davis and Bjorkman, 1988; Natarajan et al., 2016). The TCR-CD3 complex is composed of an αβTCR heterodimer with membrane embedded C-terminal helices lacking any intracellular signaling domains, a CD3γε heterodimer, a CD3δε heterodimer and a CD3ζζ homodimer. Each CD3 possesses either 1 or 3 immunoreceptor tyrosine-based activation motifs (ITAMs) in their cytoplasmic tails, which can be phosphorylated to propagate signals in the cell interior (Kane et al., 2000). This arrangement thus requires that any communication of cognate pMHC-TCR interactions into the T cell must occur through the CD3 subunits. Previously, the stoichiometry and molecular features of the association of the constituent domains were determined by different techniques (Arechaga et al., 2010; Birnbaum et al., 2014; Dong et al., 2019; He et al., 2015; Natarajan et al., 2016) but the truncated proteins and experimental conditions used raise questions about their physiological relevance.

The anchoring core of the TCR complex is the bundle of transmembrane helices (TMs) of the TCR and the CD3 chains. Highly conserved, charged residues in the TMs and membrane-proximal tetracysteine motif are required for clustering all the TCR-CD3 complex components (Call et al., 2002; Call and Wucherpfennig, 2005; Xu et al., 2006), and interactions between extracellular regions are required for bioactive TCR-CD3γε/δε complex formation (Fernandes et al., 2012; He et al., 2015) and T cell signaling (Natarajan et al., 2016). NMR chemical shift perturbation (CSP) studies involving the extracellular components of αβTCR and CD3 subunits provided residue-specific information, but suggested different binding modes (single-sided (He et al., 2015) and double-sided (Natarajan et al., 2016)) of CD3γε/δε to the αβTCR. Indeed, the peptide linking segment between the CD3 extracellular folded domains and their corresponding TMs are long enough to accommodate either a one-sided or two-sided conformation (unpublished observation). Our prior two-sided NMR model showed that, in the inactivated T cell state, the TCR Cβ subunit interacts with CD3γε through its helix 3 and helix 4-F strand regions, whereas the TCR Cα subunit interacts with CD3δε through its F and C strand regions, thereby placing the CD3 subunits on opposite sides of the TCR. This was in general agreement with an earlier electron microscopy (EM) structure of the pMHC-TCR-CD3 complex and TCRα-CD3δε SAXS structure (Birnbaum et al., 2014). While these studies provide important clues about the composition and orientation of the TCR-CD3 complex, they did not include the native TCR-CD3 TMs (Birnbaum et al., 2014; He et al., 2015; Natarajan et al., 2016). A recent 3.7 Å single-particle, non-crystalline cryo-EM structure of the human TCR-CD3 complex included the connecting peptide linker segments and the 8-TM helix bundle (without the intracellular cytosolic regions), revealing an orientation of the CD3 subunits, as well as specific contact details between individual subunits (Dong et al., 2019). This acellular structure depicted both CD3γε and CD3δε binding TCR from the same side in a non-MHC-ligated state. The absence of the plasma membrane in this structure and the use of glutaraldehyde to crosslink the TCR-CD3 subunits, however, leaves open the possibility that some of the observations may not reflect the physiologically relevant, native cell surface conformation. In addition, T cell signaling is commonly studied in mouse model systems and to what degree this conformation differs between mouse and human is not known. Indeed, major aspects of T-cell signaling are known to differ between the mouse and human immune systems (Mestas and Hughes, 2004).

Photo-crosslinking of incorporated unnatural amino acids (UAA) is a powerful tool for studying complex protein-protein interactions, molecular mechanisms, and spatiotemporal conformational states (Coin, 2018; Coin et al., 2013) and has been used to map ligand-binding sites for multiple proteins, including G protein-coupled receptors, neurokinin-1 receptor and a human serotonin transporter (Gagnon et al., 2019; Grunbeck et al., 2011; Rannversson et al., 2016; Valentin-Hansen et al., 2014). Other photo-crosslinking studies include analysis of histone-histone interactions leading to chromatin condensations (Wilkins et al., 2014) and identification of RNA-binding sites in riboprotein complexes (Kramer et al., 2014). However, this effective technique has not yet been applied extensively to study immune receptors. Here, we report a model of the mouse *in situ* cell-surface conformation of the TCR-CD3 signaling complex using constraints obtained from site-specific photo-crosslinkers that reveals a one-sided CD3ε’-CD3γ-CD3ε-CD3δ subunit arrangement around the αβTCR. We compared the model in detail to the previously solved acellular glutaraldehyde crosslinked human TCR-CD3 cryo-EM structure (Dong et al., 2019) which revealed a similar overall arrangement but with certain differences between them, especially in the TCR-CD3 interface residues.

## Results

### The TCR-CD3 complex is amenable to UAA incorporation and photo-crosslinking

To incorporate photo-crosslinkable UAAs in response to specific codons (e.g., amber stop codon) for crosslinking TCR and CD3 subunits, orthogonal tRNA/aminoacyl-tRNA synthetase (tRNA-aaRS) were designed that incorporate UAA present in cell culture media into the nascent protein within the cell at appropriate sites (Figure 1A). Previously, we co-transfected plasmids encoding tRNA-aaRS, TCR and CD3 subunits into human embryonic kidney (HEK) 293T cells and successfully incorporated UAA photo-crosslinkers p-azido-phenylalanine (pAzpa) and H-p-Bz-Phe-OH (pBpa) site-specifically into the TCR. We demonstrated the effectiveness of this approach by probing the interaction between the TCR subunits by photo-crosslinking (Wang et al., 2014). Based on this work, in the present study, we aimed to incorporate the unnatural amino acid pAzpa into previously identified TCR-CD3 interaction sites (Beddoe et al., 2009; Dong et al., 2019; Kim et al., 2009; Kuhns and Davis, 2007; Natarajan et al., 2016) on the mouse 2B4 TCR constant regions and crosslink it to neighboring mouse CD3 subunits and vice versa by UV (360 nm) activation (Figure 1B). For expression on mammalian cells, the TCR subunits (α- and β-) and CD3 subunits (γ-, δ-, ε- and ζ-) are connected by self-cleavable 2A peptides to promote stoichiometric expression of the different subunits in the TCR-CD3 complex (Figure 1C). To facilitate detection of crosslinked subunits by Western blot, the following protein tags were added to the C-terminal ends of different subunits: TCRa: c-Myc, TCRβ: V5, CD3γ: VSV-G, CD3δ: FLAG and CD3ε: HA (Figure 1C).

**Figure 1:**
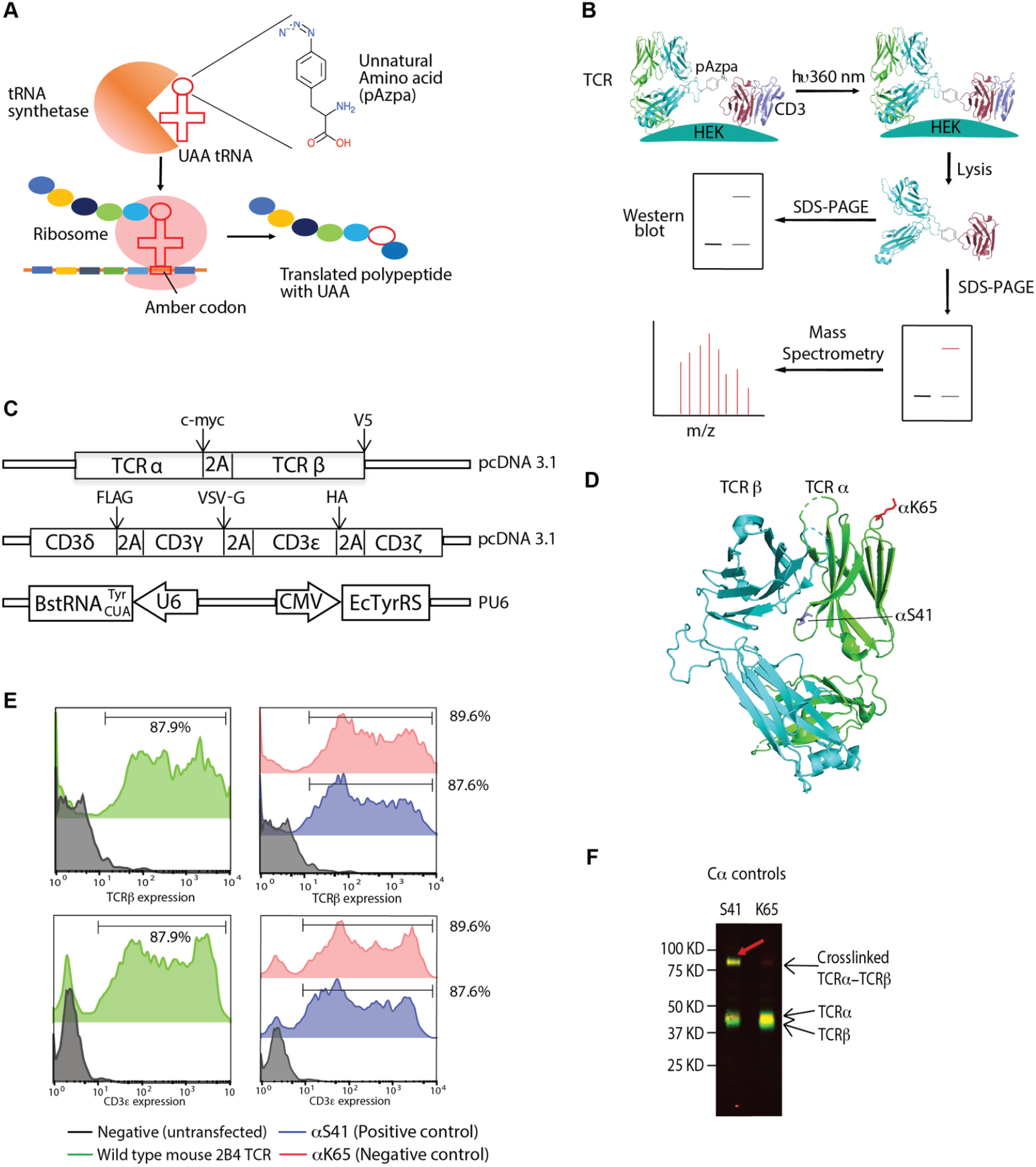
The TCR-CD3 complex is amenable to UAA incorporation and photo-crosslinking: A) Schematic overview of UAA (pAzpa) incorporation into translated protein by orthogonal tRNA/tRNA synthetase pair. B) General outline of the steps involved in the crosslinking assay in HEK293T cells. C) Illustrations of the 2B4 TCR, CD3 and tRNA-aaRS expression plasmids and locations of peptide tags utilized for Western blot identification. D) Locations of αS41 (blue), αK65 (red) in the 2B4 TCR crystal structure (PDB: 3QJF). E) TCRβ and CD3e expression profiles of wildtype 2B4 TCR (green), αS41 (blue), αK65 (red) by flow cytometry (stained with APC-conjugated H57-597 antibody – TCRβ and PE-conjugated 145-2C11 antibody – CD3ε) shows successful surface expression after UAA incorporation. The percentage of cells positive for both TCRβ and CD3ε staining is indicated. F) TCR mutant αS41 (positive control) crosslinks with TCRβ with the crosslinking band migrating between 75 to 100 kDa illustrating the feasibility of the technique to crosslink nearby subunits. The blot was stained with rabbit anti-TCRα (cMyc) antibody and mouse anti-TCRβ (V5). Anti-rabbit IRDye 680LT- and anti-mouse IRDye 800CW were used as secondary antibodies for detection.

To verify pAzpa incorporation and test photo-crosslinking, we determined the ability of TCRa with S41 and K65 mutations to crosslink TCRβ (Wang et al., 2014) (Figure 1D). The TCRa S41 mutant serves as a positive control, as it is proximal to the TCRβ subunit, and the K65 TCRa mutant serves as a negative control, as it is distal to the TCRβ subunit in the CDR region. Following transfection, 87.9% of 293T cells transfected with the wild type TCRα stained positive for both TCRβ and CD3ε, compared with 87.6% for the S41 TCRα mutation and 89.6% for the K65 TCRα mutation (Figure 1E). After UV excitation, we detected crosslinked TCRa-TCRβ for the aS41 mutant by Western blot as a band between 75 and 100 kDa, which corresponds to a size equaling TCRα+TCRβ(Figure 1F). No such band was observed for the negative control K65 TCRα mutant, and non-crosslinked TCRα and TCRβ subunits were observed at bands between 37 and 50 kDa (Figure 1F). Taken together, these results show that we can efficiently express TCR-CD3 complexes on the 293T cell surface, incorporate pAzpa into specific locations in the TCR α-subunit, and crosslink it to the adjacent TCRβ-subunit.

### CD3 subunits crosslinks with specific TCR regions indicating one-sided CD3 subunits arrangement around the TCR

Earlier studies involving mutagenesis (Kuhns and Davis, 2007), docking (Sun et al., 2004), molecular dynamics (Martinez-Martin et al., 2009), NMR (He et al., 2015; Natarajan et al., 2016), cryo-EM (Dong et al., 2019) and inference from crystal structures (Arnett et al., 2004; Kjer-Nielsen et al., 2004) identified multiple CD3 interaction sites on the TCR. We analyzed these proposed interaction sites, namely the AB loop, DE loop of TCR Cα and CC’ loop, FG loop, G strand, helix 3 and helix 4-F strand of TCR Cβ (Figure 2A, Figure 2 – table supplement 1), by UAA (pAzpa) incorporation and crosslinking, to provide a detailed model of native TCR-CD3 complex assembly *in situ* on mammalian cells.

**Figure 2:**
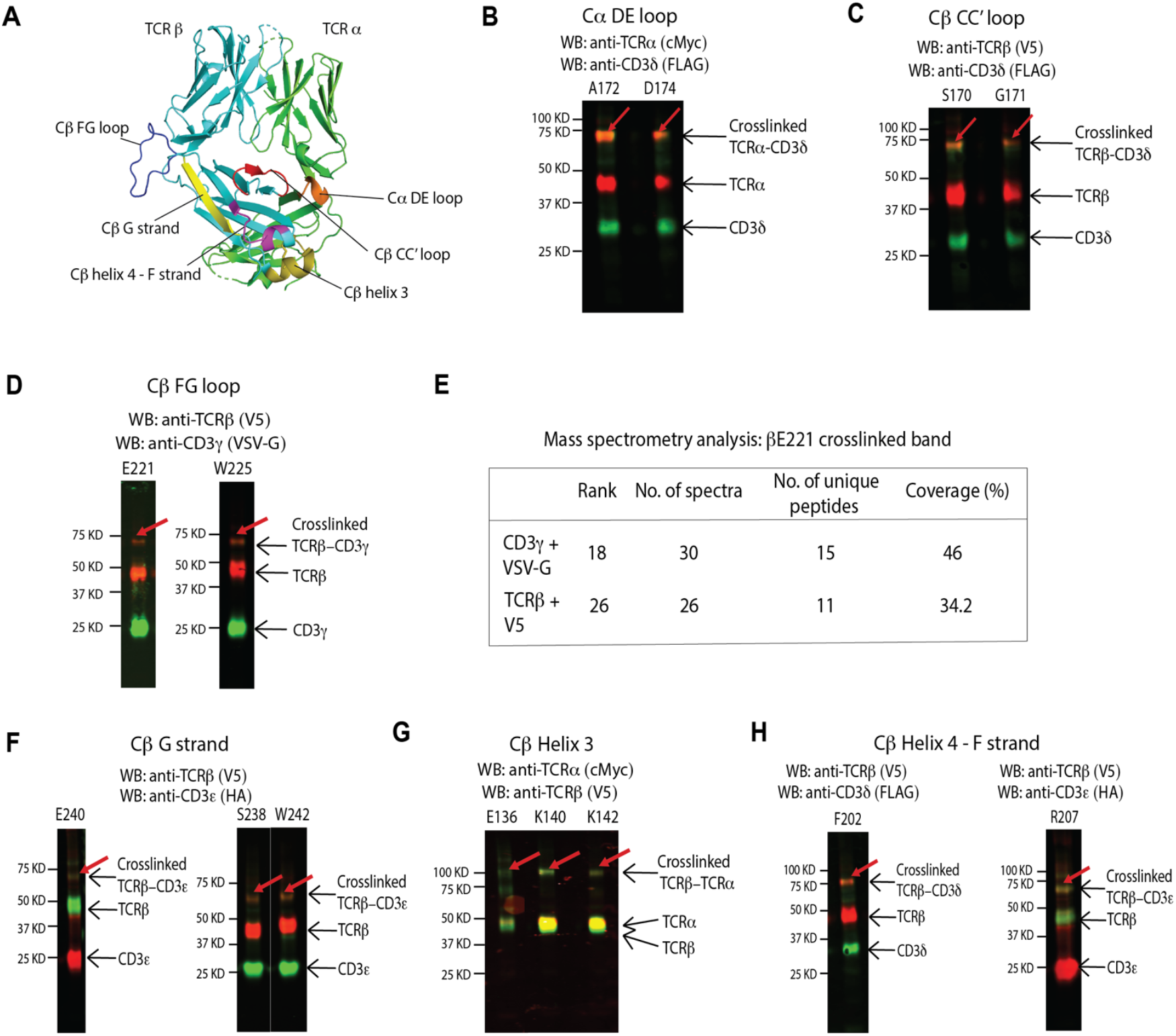
CD3 subunits crosslinks with specific TCR regions indicating one-sided CD3 subunits arrangement around the TCR. A) Location of Cα DE loop (orange), Cβ CC’ loop (red), Cβ FG loop (blue), Cβ G strand (yellow), Cβ helix 3 (olive) and Cβ helix 4 – F strand (magenta) on the 2B4 TCR crystal structure (PDB: 3QJF). B) Cα DE loop A172 and D174 crosslinks with CD3δ. The blot was stained with rabbit anti-TCRα (cMyc) antibody and mouse anti-CD3δ (FLAG). C) Cβ CC’ loop S170 and G171 crosslinks with CD3δ. The blot was stained with anti-TCRβ (V5) antibody and anti-CD3δ (FLAG). D) C βFG loop E221 and W225 crosslink to the CD3γ subunit. The blot was stained with rabbit anti-TCR β (V5) antibody and mouse anti-CD3γ (VSVG). (E) Summary of the mass spectrometry analysis on the resected Cβ E221 crosslinking band, which reveals the presence of unique CD3γ and TCRβ peptides. G) Cβ G strand S238, E240 and W242 crosslink to CD3ε. The β E240 blot was stained with mouse anti-TCRβ(V5) antibody and rabbit anti-CD3ε(HA). The β S238 and β W242 blots were stained with mouse anti-CD3ε (HA) and rabbit anti-TCRβ (V5). G) Cβ helix 3 E136, K140 and K142 crosslink to TCRα. The blots were stained with mouse anti-TCRβ (V5) and rabbit anti-TCRα (cMyc). H) Cβ helix 4 – F strand F202 and R207 crosslink with CD3δ and CD3ε, respectively. The F202 blot was stained with mouse anti-CD3δ (FLAG) and rabbit anti-TCRβ (V5). The R207 blot was stained with rabbit anti-CD3ε (HA) and mouse anti-TCRβ (V5). The crosslinking bands in each case is apparent below 75 kDa. Anti-rabbit IRDye 680LT- and anti-mouse IRDye 800CW were used for all blots as secondary antibodies for detection.

Mutagenesis of the TCR demonstrated that the TCR Cβ CC’ loop interacts with CD3εγ subunits and the Cα DE loop interacts with CD3εδ (Kuhns and Davis, 2007). To test these interaction sites, we transfected TCRs containing the following mutants into 293T cells: A172, D174 in the Cα DE loop and N164, K166, V168, S170 and G171 in the Cβ CC’ loop (Figure 2A). 87.8% of cells transfected with the A172 mutant and 58.5% of cells transfected with the D174 mutation stained positive for TCRβ and CD3ε (Figure 2 - figure supplement 1A). However, the percentage of cells that expressed TCRβ and CD3ε for the Cβ CC’ loop mutants ranged from 20.8% for the N164 mutation to 91.2% for the S170 mutation (Figure 2 - figure supplement 2A), suggesting that some of these residues might be important for the stability of the complex. Our crosslinking studies revealed that A172, D174 (both Cα DE loop), S170 and G171 (both Cβ CC’ loop), crosslinked with the CD3δ subunit of the CD3δε heterodimer (Figure 2B, 2C, Figure 2 - figure supplement 1B, 2B). We observed a crosslinked TCRα-CD3δ band for A172 and D174 and a crosslinked TCRβ-CD3δ band for S170 and G171 below the 75 KDa molecular marker (Figure 2B, 2C) corresponding to the size equaling the two crosslinked subunits. Cα 172, Cα D174, Cβ S170 and Cβ G171 cluster together in the mouse/human TCR crystal 3D structural model (Figure 2A). In the recent cryo-EM structure, the residue corresponding to A172 in the Cα DE loop, S_186_ in the human TCR Cα, contacts CD3δ in the complex. (Note: the alternate residue numbering in human cryo-EM structure is shown in smaller font size). There is no contact between the Cβ CC’ loop and CD3δ but the other end of the CC’ loop (G_182_) interacts with CD3γ in the cryo-EM structure. There is no evidence of the Cβ CC’ loop crosslinking with CD3γ in our crosslinking analysis. Overall, the relative positioning of TCRα to CD3δ determined from our crosslinking data suggests close correlation to the cryo-EM structure except for the lack of interaction between Cβ CC’ loop and CD3γ.

High-resolution X-ray structural studies and fluorescence-based experiments have demonstrated a large conformational change in the AB loop of the TCR Cα domain upon agonist binding leading to T cell activation. Moreover, deletion of the AB loop impaired T cell activation indicated by low CD69 upregulation (Beddoe et al., 2009). Allosteric changes upon antigen binding were observed in the AB loop in an NMR CSP study (Rangarajan et al., 2018). These findings led to a hypothesis that the AB loop was a possible CD3 interaction site. Based on this, we transfected and UV-crosslinked the following Cα AB loop mutants: D132 (transfection efficiency - 54.5%), R134 (23.2%) and Q136 (45.3%) (Figure 2 - figure supplement 3A, 3B). However, we observed no crosslinks between the CαAB loop and any of the CD3 subunits (Figure 2 - figure supplement 3C), suggesting that the CD3 subunits are not near the Cα AB loop. Our finding is in agreement with the cryo-EM structure (Dong et al., 2019), which showed no interactions between Cα AB loop and CD3 subunits.

Upon antigen ligation, the TCR behaves as an anisotropic mechanosensor, wherein the Cβ FG loop interacts with neighboring CD3γε by acting as a lever (Kim et al., 2009; Kim et al., 2010). Other studies using single molecule analyses, NMR and molecular dynamics revealed that the FG loop allosterically controls TCR CDR catch-bond formation, peptide discrimination and CD3 communication (Das et al., 2015; Rangarajan et al., 2018). Moreover, the FG loop is also theorized to be involved in thymic selection and T cell function and development (Sasada et al., 2002; Touma et al., 2006). Based on this and the known importance of the FG loop to T cell function, we transfected, and analyzed by crosslinking, the following TCR Cβ FG loop mutants (transfection efficiency in brackets): L219 (53.2%), E221 (84.5%), D223 (73.9%), W225 (79.0%), S229 (61.7%) and K231 (37.3%) (Figure 2A, Figure 2 – figure supplement 4A). Of these mutants, we found that Cβ FG loop residues E221 and W225 crosslink with CD3γ of the TCR-CD3 complex based on the presence of a band below the 75 kDa molecular marker indicating crosslinked TCRβ-CD3γ in the Western blot analysis (Figure 2D, Figure 2 – figure supplement 4B). However, in the cryo-EM structure CD3ε’ (belonging to CD3γε heterodimer) is in closer proximity to the CβFG loop (Dong et al., 2019). The crosslinking assay clearly indicates that CD3γ is closer to the Cβ FG loop than CD3ε, possibly due to differing surface charges between mouse and human species (see computational docking results section). To further confirm that the CD3γ subunit is closer to the Cβ FG loop than CD3ε we performed mass spectrometry analysis on the resected crosslinked band below 75 kDa from the SDS-PAGE gel for the E221 TCR Cβ FG loop mutant. More unique peptide fragments belonging to CD3γ and TCRβ were identified in the crosslinked band (15 and 11, respectively) than other TCR-CD3 complex subunits such as CD3δ (6), CD3ζ (7), TCRα (4) and CD3δ (1) (Figure 2E, Figure 2 – data supplement 1), further strengthening the possibility that CD3γ is closer to the Cβ FG loop in the cell surface conformation.

Next, we identified residues belonging to the Cβ G strand region that interact with different CD3 subunits, as it was implicated in CD3ε binding in the cryo-EM structure (Dong et al., 2019). We transfected and UV-crosslinked the following Cβ G strand mutants (transfection efficiency in brackets): N236 (57.0%), S238 (48.6%), E240 (60.0%), W242 (66.2%) and R244 (38.1%) (Figure 2A, Figure 2 – figure supplement 5A). We found that residues S238, E240 and W242 crosslinked with CD3ε, indicated by the presence of a band below the 75 kDa molecular marker in the Western blot analysis (Figure 2F, Figure 2 – figure supplement 5B). Interestingly, the residue corresponding to W242 in the cryo-EM structure, W259, interacts with CD3ε (Dong et al., 2019), indicating consistency in this aspect between our crosslinking and the cryo-EM data (Dong et al., 2019). Overall, these crosslinking experiments show that CD3γε binds to the TCR region around the Cβ FG loop and the Cβ G strand with CD3γ nearer to the FG loop.

Our earlier NMR analysis of TCR-CD3 ectodomains indicated interactions between TCR Cβ helix 3, helix 4 – F strand regions and CD3γε (Natarajan et al., 2016). This is supported by other NMR CSP studies that suggested helix 3, helix 4 regions as CD3 binding regions (He et al., 2015) and that these regions undergo allosteric changes upon antigen ligation (Natarajan et al., 2017; Rangarajan et al., 2018). Another study showed that amino acid changes in the helix 3 region led to improved TCR expression and CD3 pairing (Sommermeyer and Uckert, 2010). Based on these earlier reports, we transfected and UV-crosslinked the following mutants in 293T cells: In Cβ helix 3: E136 (63.4%), I137 (55.1%), A138 (26.8%), K140 (46.8%), Q141 (72.1%) and K142 (40.9%) (Figure 2A, Figure 2 – figure supplement 6A); in Cβ helix 4-F strand region: F202 (78.6%), H204 (84.3%), P206 (68.0%), R207 (77.9%), N208 (74.3%) and F210 (91.8%), (Figure 2A, Figure 2 – figure supplement 7A). We did not identify any crosslinks between TCR Cβ helix 3 residues and any CD3 subunits, but instead helix 3 residues E136, K140 and K142 crosslinked with the neighboring TCR α-subunit (Figure 2G, Figure 2 – figure supplement 6B). Likewise, the cryo-EM structure did not show any interactions between helix 3 residues and CD3 subunits. For the TCR Cβhelix 4-F strand mutants, crosslinking bands were observed for F202 and R207 below the 75 kDa molecular marker in the Western blot that corresponded with TCRβ+ CD3δ and TCRβ+CD3ε, respectively (Figure 2H, Figure 2 – figure supplement 7B). This is in contrast to the cryo-EM structure (Dong et al., 2019) as F strand residue H226, which is near R207, stacks against CD3γ in the cryo-EM structure (Dong et al., 2019), possibly due to differences in mouse-human surface charges. Overall, our crosslinking analysis suggests that residues F202 and R207 in the TCR Cβ helix 4 – F strand region could potentially interact with CD3δ and CD3ε, respectively, and that residues in Cβ helix 3 are not directly involved in CD3 interactions.

Overall, by incorporating UAA into different aβ TCR sites, we identified that in the TCR-CD3 complex, CD3δ is nearer in space to the Cα DE loop and Cβ CC’ loop; CD3δε is nearer in space to the Cβ helix 4-F strand regions; CD3γ is nearer in space to the Cβ FG loop and CD3ε is nearer in space to the Cβ G strand in the αβTCR-CD3 complex.

### Crosslinking identifies the sides of CD3 subunits facing the TCR in the TCR-CD3 complex

After identifying specific TCR residues that are closer to CD3 subunits, we performed reciprocal experiments to identify CD3 residues that are closer to TCR residues using the same UAA incorporation and photo-crosslinking approach. The CD3γ and CD3δ residues for UAA incorporation were selected based on their presence in the interface, near the interface or away from the interface in the human TCR-CD3 cryo-EM complex structure (Dong et al., 2019). Mutations were not introduced into the CD3ε subunit as it would not be possible to distinguish between CD3ε subunit belonging to CD3δε or CD3γε. Based on cryo-EM structure, we transfected and UV-crosslinked the following CD3δ mutants in 293T cells: In A strand: T5 (66.7%); AB loop: E8 (68.5%), D9 (71.7%); BC loop: T17 (76.3%); CD loop: V26 (31%); E strand: T35 (73.6%); and EF loop: K40 (62.8%) (Figure 3A, Figure 3 – figure supplement 1A, Figure 3 – table supplement 1). Of these mutants, we found that T5 crosslinks to TCRβ(Figure 3B, Figure 3 – figure supplement 1B) and T35 and K40 crosslinks to TCRα (Figure 3C, 3D, Figure 3 – figure supplement 1B). The conserved residue K40 (K62 in the cryo-EM structure) that crosslinks to TCRα is involved in H-bond interaction with TCRα connecting peptide residue K234 in the cryo-EM structure (Dong et al., 2019).

**Figure 3:**
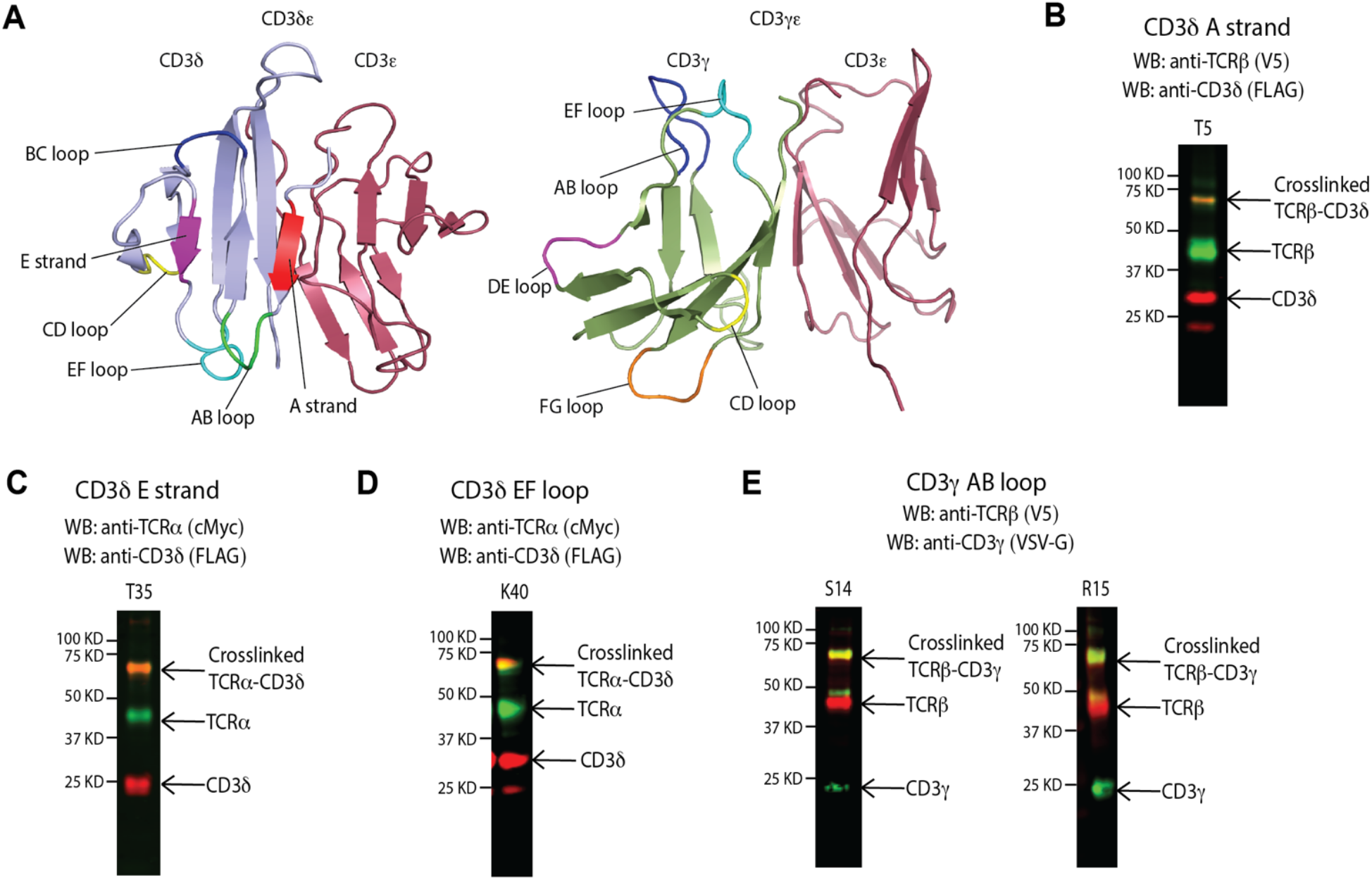
Crosslinking identifies the sides of CD3 subunits facing the TCR in the TCR-CD3 complex. A) Left, location of CD3δ A strand (red), AB loop (green), BC loop (blue), CD loop (yellow), E strand (magenta) and EF loop (cyan) on the CD3δε mouse structure. Right, location of CD3γ AB loop (blue), CD loop (yellow), DE loop (magenta), EF loop (cyan) and FG loop (orange). B) CD3δ A strand T5 crosslinks to TCRβ. The blot was stained with rabbit anti-CD3δ(FLAG) and mouse anti-TCRβ(V5). C) CD3δ E stand T35 crosslinks to TCRα. The blot was stained with rabbit anti-CD3δ(FLAG) and mouse anti-TCRα (cMyc). D) CD3δ EF loop K40 crosslinks to TCRα. The blot was stained with rabbit anti-CD3δ (FLAG) and mouse anti-TCRα (cMyc). E) CD3γ AB loop S14 and R15 crosslink to TCRβ. The blot was stained with mouse anti-CD3γ (VSV-G) and rabbit anti-TCRβ (V5). The crosslinking bands in each case is apparent below 75 kDa. Anti-rabbit IRDye 680LT- and anti-mouse IRDye 800CW were used for all blots as secondary antibodies for detection.

Residues T5 and T35 (corresponding to E27 and R57 in cryo-EM structure) that crosslink to TCRβ and TCRα, respectively, are involved in polar interactions with TCRα residue R_185_ in the cryo-EM structure. Interestingly, we do not see crosslinks for the conserved AB loop residues E8 and D9, which interact with TCRα in the cryo-EM structure (Dong et al., 2019). These differences could arise due to mouse/human species-specific surface charge differences between crosslinking and cryo-EM experiments.

For CD3γ, we transfected and UV-crosslinked the following mutants in 293T cells: AB loop: S14 (57.8%), R15 (66.8%); CD loop: D36 (59.6%); DE loop: T46 (67.6%), K47 (61.5%); EF loop: K57 (64.8%) and FG loop: A68 (63.2%) (Fig. 3A, Figure 3 – figure supplement 2A). From these, we found that AB loop residues S14 and R15 crosslink with TCRβ (Fig. 3E, Figure 3 – figure supplement 2B). The corresponding residues in the cryo-EM structure, Y_36_ and Q_37_, contact H_226_ and G_182_, respectively (Dong et al., 2019). The other mutants tested were on loops away from the AB loop, suggesting that the region around the AB loop is the one facing and nearer to TCRβ.

### CD3-crosslinking interacting Cβ G strand residues are important for T cell functionality

Based on our crosslinking results, we used site-directed mutational and functional assays in T cell hybridoma 58-/- (Letourneur and Malissen, 1989; Zhong et al., 2010) (expresses CD3 subunits but not TCRαβ) to determine whether the CD3-crosslinking TCR residues are functionally relevant to T cell activation. Multiple alanine mutations were introduced in the 2B4 TCR residues that were shown to crosslink with CD3 and mutant T cell clones were obtained by retroviral transduction (Natarajan et al., 2016; Zhong et al., 2010). Specific target sites included CβCC’ loop, CβFG loop, CβG strand and Cα DE loop (Figure 4A, 4B). Cells were co-cultured with MHCII IE^k^-expressing CHO cells (CHO-IE^k^) loaded with K5 peptide and assessed for IL-2 production via ELISA sandwich assay (Malecek et al., 2014; Natarajan et al., 2016). Activated T cell hybridoma clones containing alanine mutations at NMR identified CD3 interaction sites such as Cβhelix 3 (E136A/I137A, N139A/K140A and Q141A/K142A), Cβhelix 4-F strand (H204A/N205A, R207A/N208A and N208A/H209A) showed less than 50% IL-2 production of the wild type TCR, indicating their importance in T cell activation (Natarajan et al., 2016).

**Figure 4:**
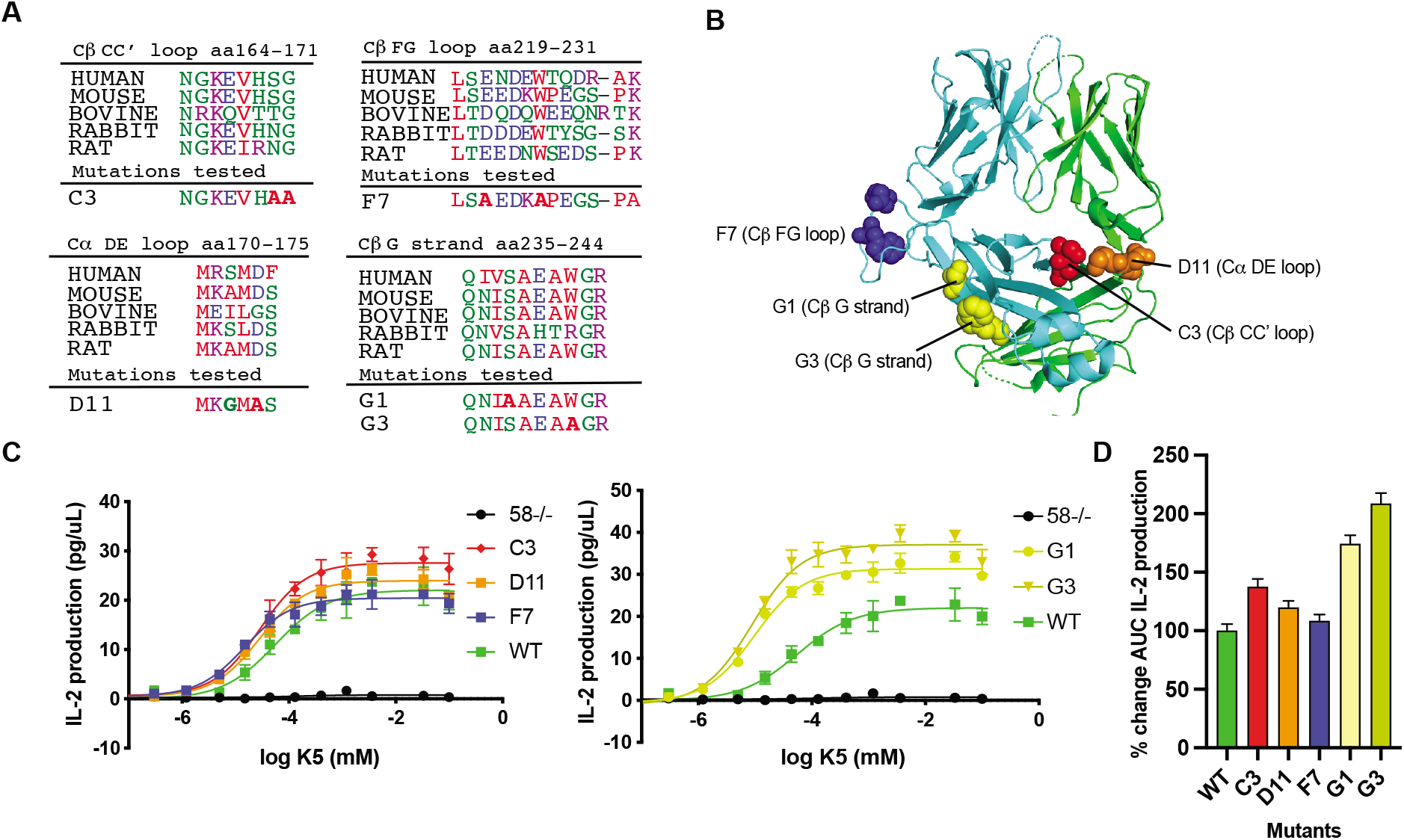
CD3-crosslinking Cβ G strand residues enhance T cell functionality. A) Sequence alignment of TCR constant regions from different species to indicate the conserved residues and the location of the TCR mutations based on cross-linking and tested in functional analysis when expressed in 58-/- T cell hybridoma. B) Locations of the mutated residues indicated on the 2B4 crystal structure. C3(S170A/G171A, red) is located in the CβCC’ loop, F7 (E221A/W225A, blue) is located in the CβFG loop, G1 (S238A, yellow) and G3 (W242A, yellow) is located in CβG strand and D11 (A172G/G174A, orange) is located in CαDE loop. C) ELISA assays (plot of IL-2 produced vs concentration of activating peptide) for mutant 2B4 T cell hybridoma clones activated with CHO/I-E^k^/K5. D) Percentage change in the area under the curve for IL-2 production between the indicated mutant T cell and wild type 2B4 T cell when activated with CHO cells expressing the cognate pMHC IE^k^/K5.

Alanine mutations at CβG strand residues S238 (G1) and W242 (G3), which crosslink to CD3ε, resulted in IL-2 production increase of 74.2 ± 7.4% and 108.6 ± 8.9% that of wild type, respectively, possibly due to mechanistic changes in the complex as a result of alanine substitutions even though S238A and W242A mutants showed surface expression comparable to the wild type (Figure 4 – figure supplement 1). Interestingly, S238R and W242R mutations increased TCR β-subunit surface expression more than 150% compared to wild type in J.RT3-T3.5 cells (Fernandes et al., 2012) indicating S238 and W242 play vital role in complex assembly and possibly T cell signaling. Overall, based on activation assays, CβG strand residues S238 and W242 play a crucial role in T cell signaling, similar to Cβ helix 3 and helix 4-F strand residues (Natarajan et al., 2016). However, the functional effects of the crosslinking mutants were not as intense as the NMR-identified mutants (Natarajan et al., 2016), CβS170A/G171A (C3, CC’ loop), Cβ E221A/W225A (F7, FG loop) and Cα A172G/D174A (D11, DE loop), which led to changes in T cell activation of +37.6 ± 6.6%, +8.5 ± 5.4% and +20 ± 5.5%, respectively (Figure 4C, 4D).

### Computational docking reveals a CD3ε’-CD3γ-CD3ε-CD3δ model for CD3 binding

To generate a crosslink-guided 3D model of the TCR-CD3 complex using crosslinking constraints (Figure 5A), we used computational molecular protein-protein docking (Fernandez-Recio et al., 2003) to generate all possible unclashed, compact conformers of mouse CD3γε and CD3δε domains with the mouse 2B4 TCR domains (Figure 5 – table supplement 1). The mouse TCR-CD3 components were modeled from available structures of human proteins (Figure 5 – table supplement 1). Thousands of unclashed, compact TCR-CD3γε-CD3δε extracellular conformations were ranked based on the following hierarchy of constraints: 1) geometric and spatial compatibility of the C-termini of the CD3γε and CD3δε extracellular folded domains with the N-termini of their corresponding TM helices in the cryo-EM TM bundle, 2) CD3 subunit crosslinks to TCR chains, 3) TCR crosslinks to CD3 subunits, 4) calculated biophysical energy (van der Waals, solvation electrostatics, hydrogen bonding), 5) absence of encroachment on plasma membrane location, 6) absence of clash with 2C11 antibody structure bound CD3ε and 7) absence of encroachment on pMHC binding site.

**Figure 5:**
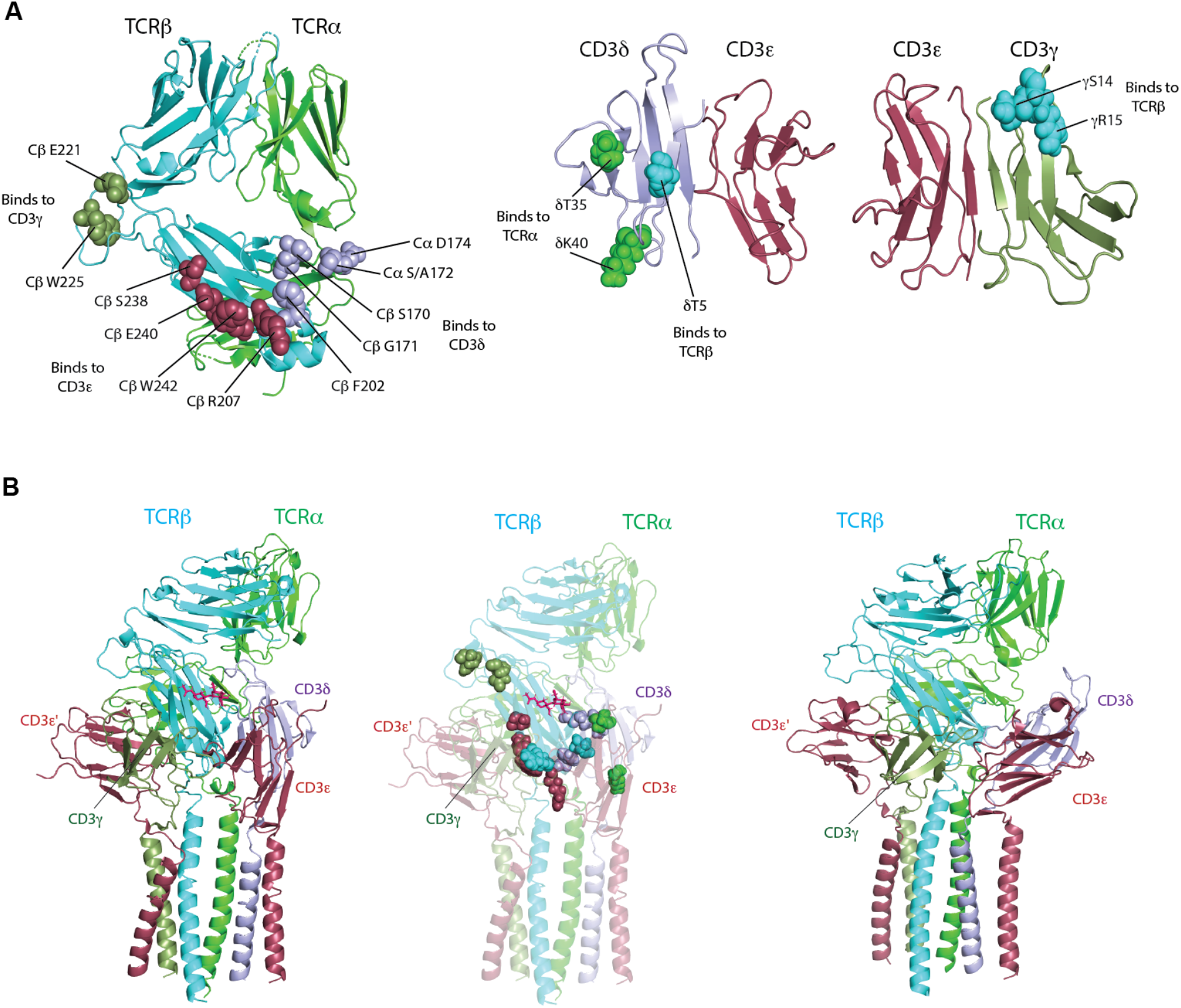
Computational docking reveals a CD3ε’-CD3γ-CD3ε-CD3δ model for CD3 binding. A) Left, crosslinking TCR residues indicated as spheres on the crystal structure of 2B4 TCR (with human constant domains, PDB: 3QJF). Residues interacting with CD3γ indicated in smudge green, with CD3ε indicated in raspberry and with CD3δ indicated in light blue. Center, crosslinking CD3δ residues indicated as spheres on the mouse CD3δε structure. Residues interacting with TCRα indicated in green, and with TCRβ indicated in cyan. Right, crosslinking CD3γ residues indicated as spheres on the mouse CD3γε structure. Residues interacting with TCRβ are indicated in cyan B) Left, docked TCR-CD3 complex structure based on crosslinking derived constraints. CD3γε interact primarily on the TCRβ face of the complex. CD3δε interacts in a region involving the interface of TCRα-TCRβ. The transmembrane bundle is derived from the cryo-EM human TCR-CD3 transmembrane helical bundle (PDB: 6JXR) (Dong et al., 2019). Center, TCR-CD3 crosslink-guided model depicted in cartoon representation (60% transparency) with crosslinking

TCR-CD3 residues in spheres. Same color scheme for the residues in spheres as in A). Right, Cryo-EM TCR-CD3 structure. TCRα, TCRβ, CD3γ, CD3δ, CD3ε/ε’ are indicated in green, cyan, smudge, light blue and raspberry, respectively.

Based on these criteria, the top scoring TCR-CD3 conformation that satisfies all crosslinking constraints (Figure 5 – table supplement 2) is similar to the cryo-EM structure (Figure 5B, left, right). The TMs of all the subunits from the cryo-EM structure and connecting linkers were modeled into the crosslink-guided model and found to be consistent with the model (e.g. the CD3 domains were docked without constraints and the cross-links were recorded without bias: if the TM helices clashed with the CD3 domains, this would be an indication of either an invalid docking method or invalid cross-links, as would the inability of the linkers to connect the CD3 C-termini to the TM N-termini). This final structural model had an overall contact area of 491.2 Å^2^ between crosslinked TCR residues and CD3γε and a contact area of 342.4 Å^2^ between crosslinked TCR residues and CD3δε, with energies of −23.6 and −14.5, respectively. The CD3ε’ chain of the CD3γε’ heterodimer is behind the Cβ FG loop (Figure 5B, left) rather than directly below the FG loop, as seen in the cryo-EM structure (Figure 5B, right). The crosslink-guided model shows that the CD3γ subunit is closer to the Cβ FG loop, as we detected crosslinking between the Cβ FG loop residues and CD3γ (Figure 5B, center, Figure 5 – figure supplement 2). The electrostatic surfaces participating in binding interfaces in the individual components of the human TCR-CD3 complex differed when compared to their modeled mouse counterparts, especially on the CD3 subunits (Figure 5 – figure supplement 1), suggesting that there could be differences between the human and mouse TCR-CD3 complexes, as seen in our model, even though the overall arrangement of the components are similar. The distances between the center of mass of the CD3δε and CD3γε relative to the TCR when compared to the cryo-EM structure are 14.7 and 36.05, respectively. The location of CD3δε on the crosslink-guided TCR-CD3 structure that was implicated in CD3δ binding is nearly identical as the same regions (A172 in crosslinking and S_186_ in cryo-EM, both belonging to Cα DE loop) in the cryo-EM structure, although the relative orientation of CD3δε in crosslink-guided structure is different than the cryoEM structure (Figure 5 – figure supplement 2). Overall, our crosslinking analysis combined with computational docking identifies a cell-surface native conformation of the aβTCR-CD3 complex that is overall similar to the human cryo-EM structure but with differences in interface contacts. Our structure identifies a CD3ε’-CD3γ-CD3ε-CD3δ model for CD3 binding with contacts between CD3γ (belonging to CD3γε) and CD3ε (belonging to CD3δε) (Figure 5B, Figure 5 – figure supplement 2).

## Discussion

A molecular phenomenon that can confound precise translation of molecular observations to clinical relevance is structural/atomic accuracy of 3D receptor models. X-ray crystallography, NMR and cryo-EM are the leading structural biology techniques that provide detailed and tangible atomistic details about protein structure and interactions. However, some biologically relevant conformations may not be identified via these techniques due to experimental and nonnative conditions used. Site-specific crosslinking using unnatural amino acids combined with computational analysis can provide a robust alternative towards obtaining both atomistic and species-specific information on intermediate or dynamic states with small amounts of protein and, significantly, on *in situ* states present in complex cellular environment (Coin, 2018; Grunbeck et al., 2011; Valentin-Hansen et al., 2014). As far as we know, our study is the first to successfully photo-crosslink and subsequently provide a native structural model of an immune protein receptor complex. The results from our study indicating overall similarity between cryo-EM structure and crosslinking model validate photo-crosslinking-docking technique as an attractive option for structural/*in situ* analysis of protein complexes. A similar approach can be undertaken to study other immune protein complexes in their native states, such as γδ TCR-CD3 complex, B cell receptor complex, CD19/CD21 coreceptor complex.

For unnatural amino acid (pAzpa) incorporation and crosslinking, we co-transfected tRNA-aaRS and mouse TCR and CD3 plasmids (with amber mutations in specific sites on TCR or CD3) into 293T cells, and expressed and UV crosslinked 47 different mutants. From this, 16 specific TCR-CD3 subunit crosslinks were identified, including residues in the Cα DE loop crosslinking with CD3δ, the Cβ CC’ loop crosslinking to CD3δ, the Cβ FG loop crosslinking to CD3γ, the Cβ G strand crosslinking to CD3ε and the Cβ helix 4-F strand crosslinking to CD3δ and CD3ε(Figure 5A). Similarly, the CD3δ A strand crosslinks to TCRβ, CD3δ E strand, the EF loop crosslinks to TCRα and the CD3γ AB loop crosslinks to TCRβ. Utilizing these specific crosslinks as distance restraints in a biophysical search of conformations, we visualized an *in situ* cell-surface model of the TCR-CD3 complex (Fernandez-Recio et al., 2003).

Comparing the crosslink-guided model with the recently published cryo-EM model (Dong et al., 2019), we found that the CD3 arrangement from the crosslink-guided model is largely comparable to the CD3 arrangement in the cryo-EM structure (Figure 5B). The gross locations of the CD3-TCR interfaces within the complexes are all similar between our photo-crosslinking and the cryo-EM, however, the Cβ FG loop is above and in-between CD3γ and CD3ε’ in the crosslink-guided model. This is different from the cryo-EM structure, which places FG loop above CD3ε’ in the CD3γε heterodimer. One reason for the difference could be the usage of glutaraldehyde crosslinking to fix the complex in one acellular conformation in cryo-EM analysis(Dong et al., 2019). Our crosslink-based conformer could represent the true resting conformation of the complex resulting from the physiological cell surface condition with Cβ FG loop closer to CD3γ in the human TCR-CD3 complex as well. However, examining this experimentally through photo-crosslinking-computational docking is beyond the scope of this current research. The other and most probable reason could be because of the species-specific differences in the amino acid composition of the electrostatic surfaces of the mouse proteins used in our study and the human proteins in the cryo-EM study (Figure 5 – figure supplement 1). By observing overall consistent locations of the CD3-TCR interfaces, our findings independently validate the mechanism of this aspect of TCR signaling between mice and humans, but, the regions of the CD3-TCR interaction that differ between mouse and human suggest that signaling thresholds, and therefore pharmacology, may be different between the species. Species-specific differences between mice and humans can confound translation of observations in more easily controlled experiments in mice to clinical relevance. Drug candidates targeting TCR-CD3 complex derived from pre-clinical murine models, ‘humanized’ murine models with human CD3 subunits (Crespo et al., 2021; Ueda et al., 2017) and CD3 copotentiation (Becher et al., 2020; Hoffmann et al., 2015) should take our crosslink-guided model into account before translating the pharmacology to human studies. For investigators using mouse systems to investigate TCR signaling and phenotypes, our crosslink-guided model may serve as a useful reference point for interpreting translatability of findings to the human TCR via comparison with the cryo-EM model.

Our crosslink-guided model differs substantially from our previously reported NMR-based model (Natarajan et al., 2016), which was based on CSP data that showed peak intensity losses or peak shits in the TCR upon CD3γε and CD3δε addition. These sites include Cβ helix 3, Cβ helix 4-F strand, Cβ FG loop, Cα F strand, Cα C strand and Cα tail (Natarajan et al., 2016). This spectral change could result from direct CD3 subunit interaction to the particular TCR site or from conformational changes at the site upon CD3 binding elsewhere on the TCR. A ranking mechanism similar to the one used in the crosslinking study was used for TCR-CD3γε and TCR-CD3δε docking. Based on this, our top-scoring TCR-CD3γε and TCR-CD3δε docking models showed that the CD3 heterodimers interact on opposite sides of the TCR (Natarajan et al., 2016). One possible reason for this discrepancy could be the absence of membrane, CPs and TMs in the soluble protein domains used in the NMR study, which could allow for orientation of the CD3 molecules away from the membrane, instead of proximal to the membrane. Further, IL-2 production upon activation of NMR-identified Cβ helix 3 mutants indicated a loss of >50% compared to the wild type TCR (Natarajan et al., 2016). Moreover, other studies have identified some of the same TCR sites (Cβ helix 3 and helix 4) as CD3 interaction regions and are involved in allosteric interactions upon antigen ligation (He et al., 2015; Natarajan et al., 2017; Rangarajan et al., 2018). Thus, this discrepancy remains unresolved for Cβ helix 3 and other sites such as the Cα F and C strands. Importantly, in this study, we identified Cβ G strand residues S238 and W242 as playing a role in T cell activation. Interestingly, these same residues show increased occupancy in the TCR-CD3 interface during force-based steered molecular dynamics simulations, thereby strengthening TCR-CD3 interactions under force (Z. Yuan, 2021). Thus, the G strand residues-S238 and W242, conserved between mouse and human, are possible candidates for protein engineering to enhance TCR signaling.

Based on data from earlier NMR and cryo-EM studies (Arechaga et al., 2010; Birnbaum et al., 2014; Dong et al., 2019; He et al., 2015; Natarajan et al., 2016), we inferred that the extracellular part of the TCR-CD3 complex could exist in multiple, biologically-relevant conformations on the T cell surface and, here, we sought to identify them. This kind of conformational switch is not uncommon in the TCR-CD3 transmembrane space, as the TCRα transmembrane helix exists in L- and E-states (Brazin et al., 2018), CD3ζζ juxtamembrane regions exist in open and closed conformations (Lee et al., 2015), and TCRβ switches between inactive and active conformations upon cholesterol binding and unbinding (Swamy et al., 2016). Using photo-crosslink-guided computational molecular docking we visualized a conformer that is similar overall to the recent 3.7 Å cryo-EM structure (Dong et al., 2019), providing validation of this model of the TCR-CD3 signaling complex. Extending the crosslink-guided model via an antigen activation system could reveal the broader mechanism by which pMHC activates the TCR-CD3 complex and identify structure-activity relationships that can be exploited to modulate signaling pharmaceutically, with potential benefits for the treatment of cancer, infectious diseases and autoimmune diseases.

## Materials and Methods

### Plasmid construction

The tRNA synthetase for recognition of pBpa in PU6-pBpa plasmid, a generous gift from Peter G. Schultz, Scripps Research Institute, was replaced with tRNA synthetase for pAzpa to create PU6-pAzpa plasmid(Wang et al., 2014). This PU6-pAzpa plasmid contains mutant E.coli tyrosyl-tRNA synthetase (EcTyrRS), one copy of *B. stearothermophilus* tRNAs (BstRNA) and human U6 small nuclear promoter (U6)(Wang et al., 2014). cDNA encoding mouse TCR 2B4 α (with c-Myc-tag) and β (with V5-tag) sequences with 2A sequence linking each other were cloned into pCDNA3.1/Zeo(+) vector (Life Technologies) using Not1 and Xho1 restriction enzymes. Similarly, cDNA encoding mouse CD3δ (with FLAG-tag), mouse CD3γ (with VSV-G tag), mouse CD3ε (with HA-tag) and CD3ζ interconnected with 2A sequence were cloned in pCDNA3.1/Zeo(+) vector using Not1 and Xho1 restriction enzymes. Amber (TAG) codons were introduced site-specifically in the 2B4 TCR and CD3 plasmid using Quikchange mutagenesis kit (Agilent).

### Transfections into HEK293T cells

HEK293T cells (ATCC) were cultured in DMEM media, supplemented with 10% FBS, sodium pyruvate, non-essential amino acids, glutaMAX-1, penicillin-streptomycin and β-mercaptoethanol and grown at 37 °C, 5% CO2 to 80% confluency in a collagen-coated 6-well plate before transfections. To incorporate unnatural amino acids, pAzpa (p-azido-phenylalanine), into predetermined sites on the TCR and CD3 extracellular regions (Table S1, S2), plasmid expressing amber suppressor tRNA-aminoacyl-tRNAsynthetase (tRNA-aaRS) – PU6-pAzpa was co-transfected with plasmids expressing full-lengths mutant 2B4 TCR and CD3 subunits using Xfect transfection kit (TaKaRa). For incorporating pAzpa into TCR sites, as optimized previously, 7.5 μg of TCR:CD3 in 8:1 ratio and 2.5 μg of PU6-pAzpa plasmids were co-transfected into HEK293T cells(Wang et al., 2014). For incorporating pAzpa into CD3 sites, 7.5 μg of TCR:CD3 in 4:1 ratio and 2.5 μg of PU6-pAzpa plasmids were co-transfected into HEK293T cells. After 4 hours of culture at 37 °C, 5% CO2, the media was replaced with fresh DMEM media containing 1 mM pAzpa (Chem-Implex International) and cultured for 48 hours.

### Flow cytometry analysis

After 48 hours of culture, the cells were harvested and washed in FACS buffer (PBS + 2% FBS). A small portion of the cells were treated with allophycocyanin (APC) anti-TCRβ (clone H57-597, eBioscience) and phycoerythrin (PE) anti-CD3ε (clone 145-2C11, eBioscience) in FACS buffer for 30 minutes. Subsequently, the samples were analyzed for TCRβ and CD3ε expression in FACSCalibur (BD Biosciences) and data was analyzed using FlowJo (ver 10.5.3).

### Photo-crosslinking, Immunoprecipitation and Western blotting

Cells were photo-crosslinked by exposing them to 360 nm UV light source for 45 minutes on ice. Following that, the cells were washed in antibody buffer - Hank’s balanced salt solution/2% FBS/0.05% (m/v) sodium azide. The cells were treated with 25 ug/mL biotinylated mouse anti-CD3ε(clone 145-2C11, eBioscience) for 30 minutes and washed in 1X TBS (Tris-buffered saline). The cells were lysed in TBS/1% (v/v) IGEPAL-630 (sigma) containing 1X Complete protease cocktail inhibitors (Roche). The TCR-CD3 complex was purified from the lysate using Dynabeads M-280 streptavidin (Invitrogen). The beads were subsequently washed with 1X TBS and boiled with SDS-PAGE reducing buffer with β-mercaptoethanol. The subunits were resolved by SDS-PAGE electrophoresis and transferred to nitrocellulose membranes (ThermoFisher). For Western blot analysis, the following pairs of primary antibodies were used: 1) TCRα-TCRβ crosslinking: rabbit anti-c-Myc (Genscript) and mouse anti-V5 (Genscript); 2) TCRα-CD3δ crosslinking: rabbit anti-cMyc and mouse anti-FLAG (Genscript); 3) TCRα-CD3δ crosslinking: mouse anti-cMyc (Genscript) and rabbit anti-FLAG (Genscript); 4) TCRβ-CD3δ crosslinking: mouse anti-V5 and rabbit anti-FLAG (Genscript); 5) TCRβ-CD3δcrosslinking: rabbit anti-V5 and mouse anti-FLAG; 6) TCRβ-CD3γ crosslinking: rabbit anti-V5 (Genscript) and mouse anti-VSV-G (Abcam); 7) TCRβ-CD3ε crosslinking: rabbit anti-V5 and mouse anti-HA (Genscript); 8) TCRβ-CD3εcrosslinking: mouse anti-V5 and rabbit anti-HA (Genscript); 9) TCRα-CD3γ crosslinking: rabbit anti-cMyc and mouse anti-VSV-G. The following secondary antibodies were used for detection: IRDye 680LT-conjugated donkey anti-rabbit IgG (H+L) (LI-COR) and IRDye 800CW-conjugated donkey anti-mouse IgG (H+L) (LI-COR). Images were collected using LI-COR Odyssey and analyzed using Image Studio Lite (LI-COR, ver 4.0.21).

### Functional analysis with mutant T cell hybridoma

Mouse 58-/- T cell hybridoma cells (Letourneur and Malissen, 1989), which expresses mouse CD3 but not TCRαβ, (from David Kranz, University of Illinois) and Chinese hamster ovary (CHO) cells expressing I-E^k^ (Krogsgaard et al., 2005)(from Mark M. Davis, Stanford University) were cultured in RPMI 1640 medium and DMEM respectively, supplemented with 10% FBS, sodium pyruvate, non-essential amino acids, glutaMAX-1, penicillin-streptomycin and β-mercaptoethanol. The mutant mouse 2B4 TCR constructs were generated by PCR using overlapping primers containing the mutant sequences and cloned into the pCDNA3.1 vector. Retroviral transductions of the hybridoma cells were done as described previously (Zhong et al., 2010). The transduced cells were stained with PE anti-CD3ε(clone 145-2C11) and APC anti-TCRβ(clone H57-597) antibodies. The transduced cells were sorted, expanded for 6 days, quantified for TCRβ/CD3ε expression, and prepared for the cytokine assay. 10^4^ CHO-IE^k^ cells were loaded with different concentrations of a variant of moth cytochrome c (K5) (Krogsgaard et al., 2005) peptide and incubated with 10^4^ T cell hybridoma clones (wild type and mutants) in triplicates for 16 hours at 37 °C, 5% CO2. A standard ELISA sandwich was used to quantify cytokine IL-2 production (Malecek et al., 2013). The area under the curve for wildtype and mutant IL-2 production, a cumulative response measure, was calculated after non-linear fitting using Prism (GraphPad software).

### TCR and CD3 subunit structure generation and complex docking

#### Crosslink-guided Model

3D models of individual murine CD3 domains (CD3γ, CD3δ) were built by homology modeling using ICM-Pro software (Molsoft LLC. La Jolla CA)(Fernandez-Recio et al., 2003) applied to different structures as templates found in the PDB database as shown in Table S3, as for the murine CD3ε its structure was taken from a published crystal structure of the monomer binding to the antibody OKT3 (PDB: 1SY6),. The coordinates of the TMs of all subunits relative to the TCR subunits were inherited from the human cryo-EM structure (PDB: 6JXR)(Dong et al., 2019). 3D models of the two CD3 hetero-oligomers were docked to the 3D model of the 2B4 murine TCR. Absence of clashes and compact conformations were identified by calculated, estimated free energy of the complex, as previously described (Fernandez-Recio et al., 2002a, b, 2003; Garzon et al., 2009), which includes terms for van der Waals, electrostatics, hydrogen bonding and solvation and the energy units approximate kcals in free energy calculations. All compact (all CD3 and TCR domains contacting at least one other domain) and conformations without clashes were retained and their calculated free energy score was re-weighted by their contact area of the UAA side-chains with the CD3 domains and the distance between these UAA side-chains and the CD3. Docked conformations that were inconsistent with the length of the linking segments that connect the TMs with the CD3 folded domains were discarded. Conformations impinging on the membrane or the 2C11 antibody were discarded, if there existed at least one unclashed, compact, cross-link compatible conformation that did not impinge on the membrane or 2C11.

## Acknowledgements

We thank Peter G. Schultz, Scripps Research Institute for the PU6-pBpa plasmid. We also thank David Kranz (University of Illinois) for providing us 58 -/- T-cell hybridoma and Mark M. Davis (Stanford University) for providing us with Chinese hamster ovary (CHO) cells expressing I-E^k^. We also thank Thomas P. Sakmar (Rockefeller University) for protocols and advice on photo-crosslinking technique. We thank Duane Moogk (McMaster University) and Yury Patskovsky (NYU Grossman School of Medicine) for helpful discussions and critical reading of the manuscript. We thank Eric Ni (Yale University) for helpful discussions. This work was supported by the NIH grant NIGMS R01 GM124489 (to M.K).

## Footnotes

### Author Contributions

Experiments were conceptualized by A.N., W.W., T.C. and M.K. Photo-crosslinking experiments were performed by A.N. Computational docking was performed by M.B.F. T cell activation experiments were performed by A.N. Crosslinking TCR, CD3 mutant constructs and mutant retroviral TCR constructs were generated by W.W., T.L., S.B. and H.S. Data was analyzed and interpreted by A.N., T.C., and M.K. The original draft was written by A.N. and the final draft was reviewed and edited by A.N., T.C., and M.K.

The authors declare no competing interests.

## Supplementary Figures

**Figure 2 – figure supplement 1:**
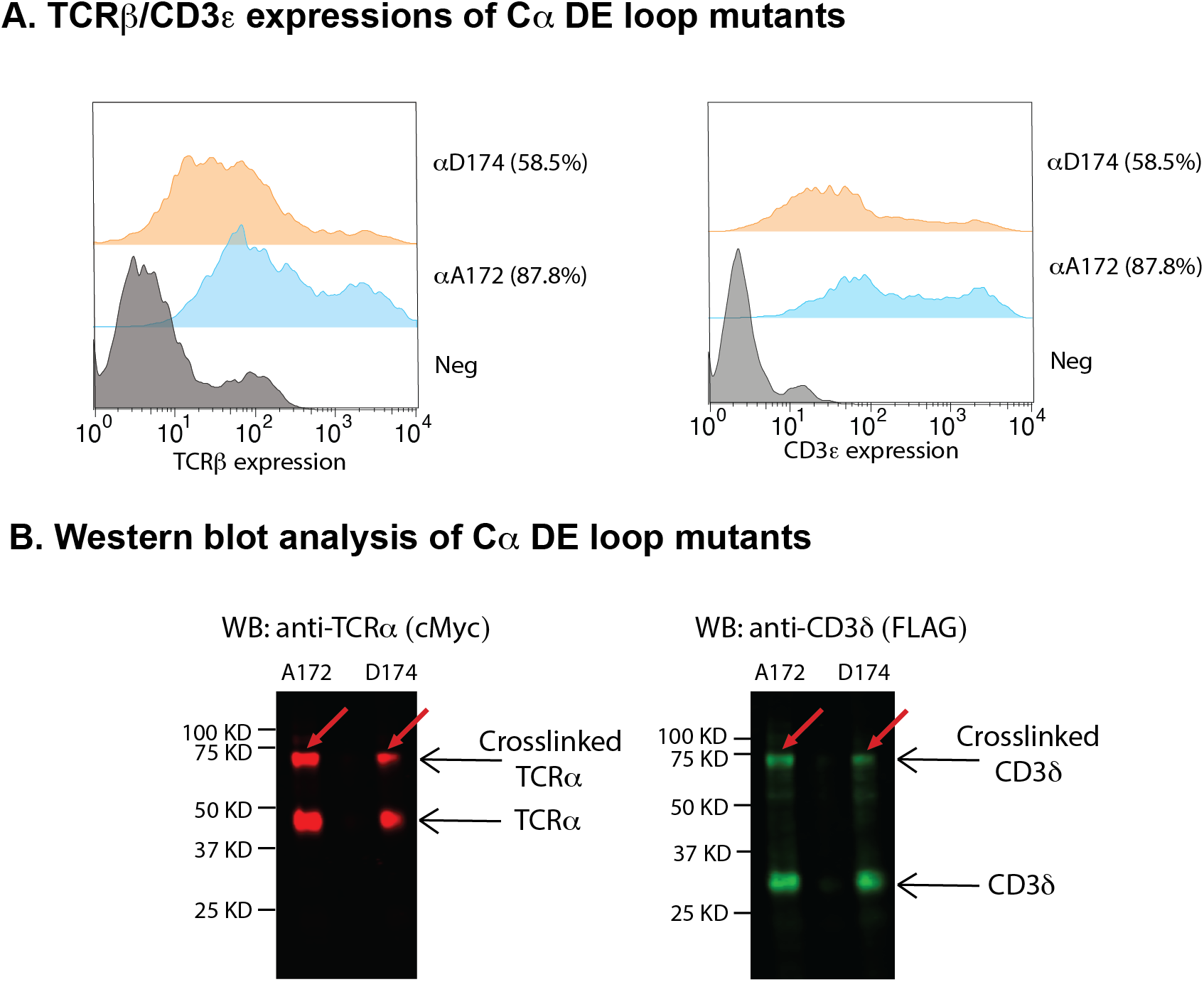
CD3δ interacts with the TCR Cα DE loop. A) TCRβ and CD3ε expression histograms of Cα DE loop mutants – αA172 and αD174 by flow cytometry. The percentage of cells positive for both TCRβ and CD3ε staining is indicated. B) Western blot analysis of Cα DE loop mutants - αA172 and αD174. CD3δ crosslinked bands for αA172 and α D174 are apparent around 75 kDa. The blot was stained with rabbit anti-TCRα (cMyc) antibody and mouse anti-CD3δ (FLAG). Anti-rabbit IRDye 680LT- and anti-mouse IRDye 800CW were used as secondary antibodies for detection.

**Figure 2 – figure supplement 2:**
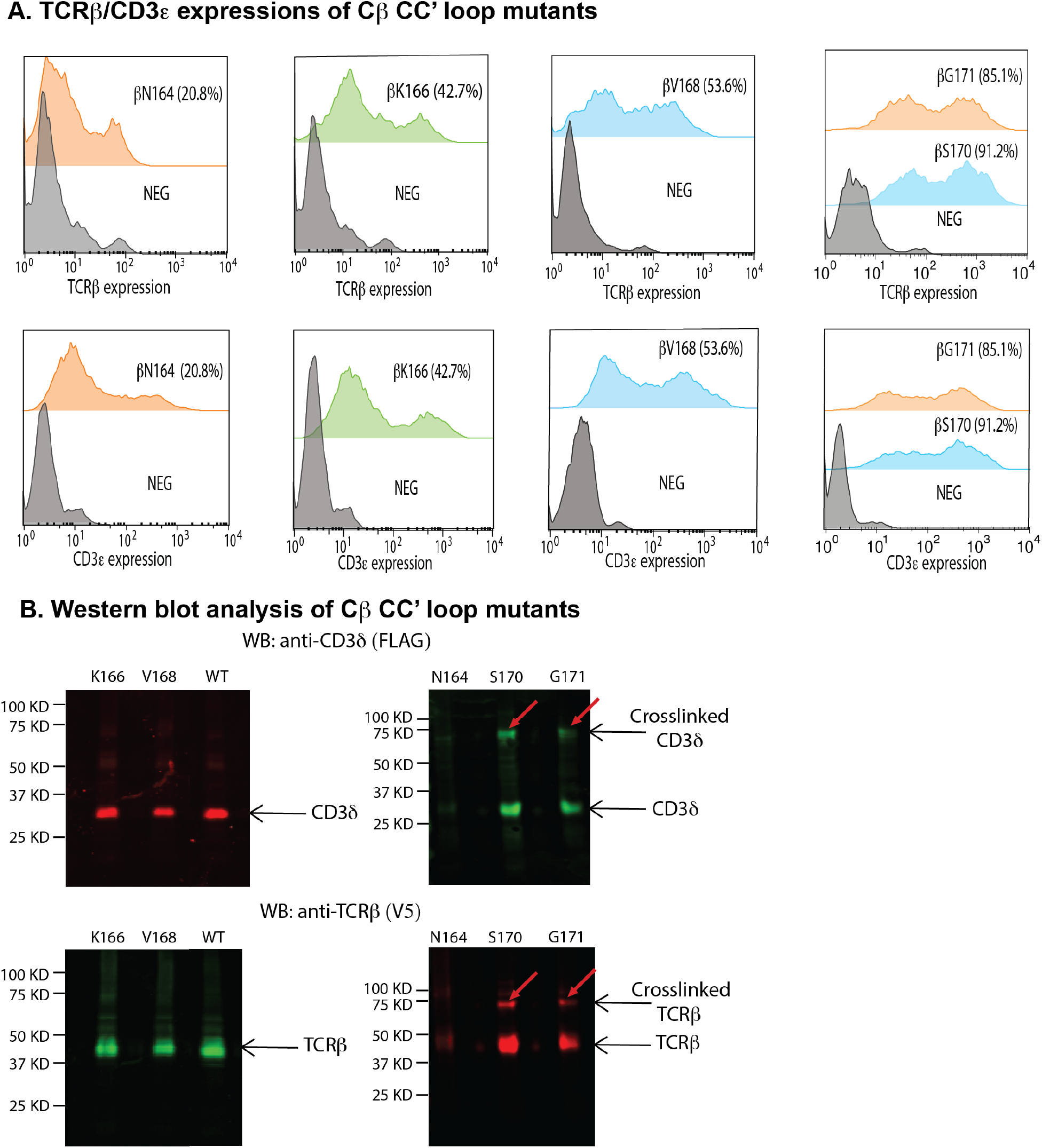
CD3δ interacts with the TCR Cβ CC’ loop. A) TCRβ and CD3ε expression histograms of Cβ CC’ loop mutants - β N164, βK166, β V168, β S170 and β G171 by flow cytometry. The percentage of cells positive for both TCRβ and CD3ε staining is indicated. B) Western blot analysis of Cβ CC’ loop mutants - βN164, βK166, βV168, βS170 and βG171. CD3δ crosslinked bands for βS170 and βG171 are apparent around 75 kDa. The blot was stained with rabbit anti-TCRβ(V5) antibody and mouse anti-CD3δ (FLAG). Anti-rabbit IRDye 680LT- and anti-mouse IRDye 800CW were used as secondary antibodies for detection.

**Figure 2 – figure supplement 3:**
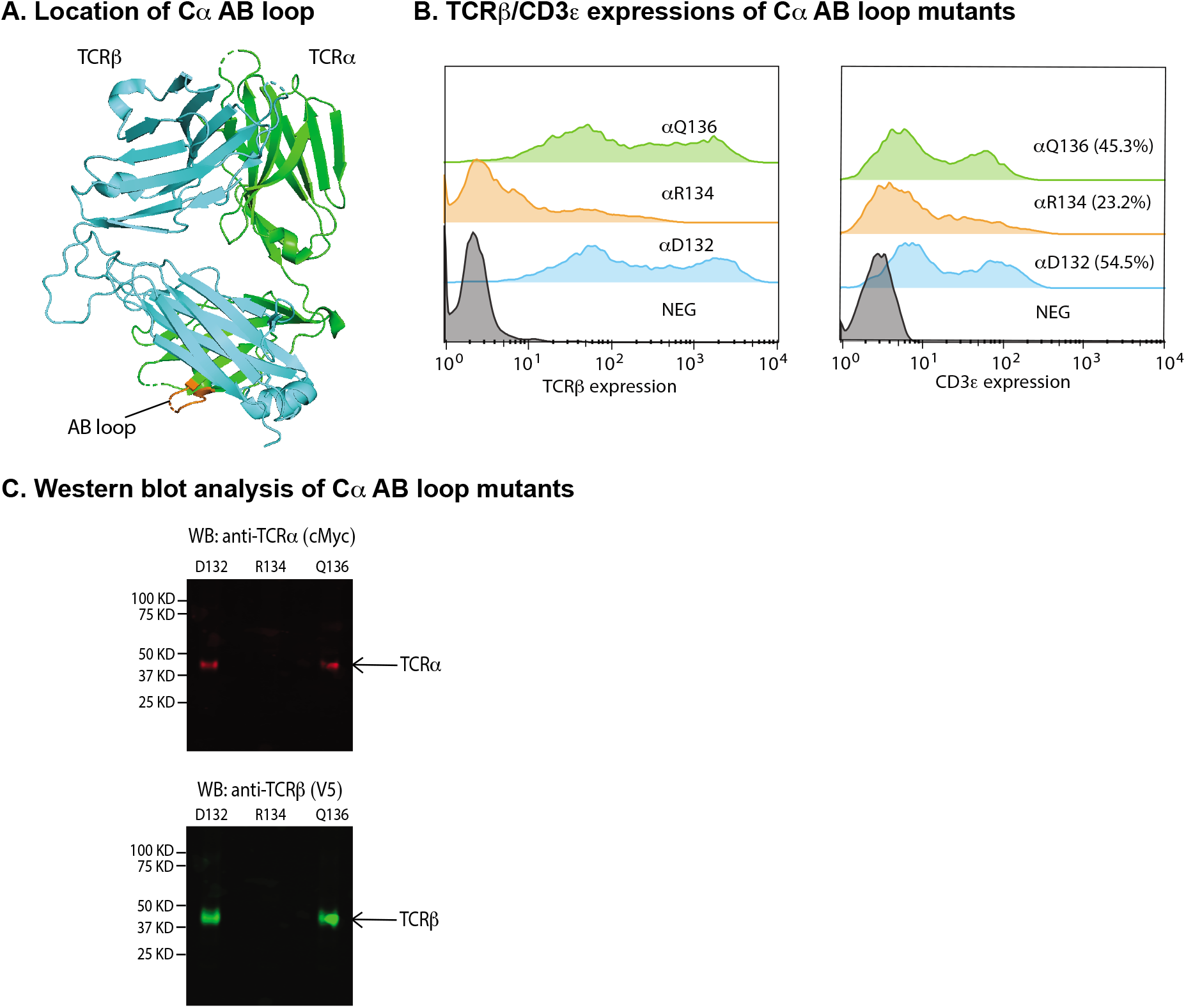
The TCR Cα AB loop is not near any CD3 subunits. A) Location of the Cα AB loop (in orange) on the 2B4 TCR crystal structure (PDB: 3QJF). B) TCRβ and CD3ε expression histograms of Cα AB loop mutants - αD132, αR134 and αQ136 by flow cytometry. The percentage of cells positive for both TCRβ and CD3ε staining is indicated. C) Western blot analysis of Cα AB loop mutants - αD132, αR134 and αQ136. No crosslinking bands were evident for these mutants. The blots were stained with rabbit anti-TCRα (cMyc) antibody and mouse anti-TCRβ (V5). Anti-rabbit IRDye 680LT- and anti-mouse IRDye 800CW were used as secondary antibodies for detection.

**Figure 2 - figure supplement 4:**
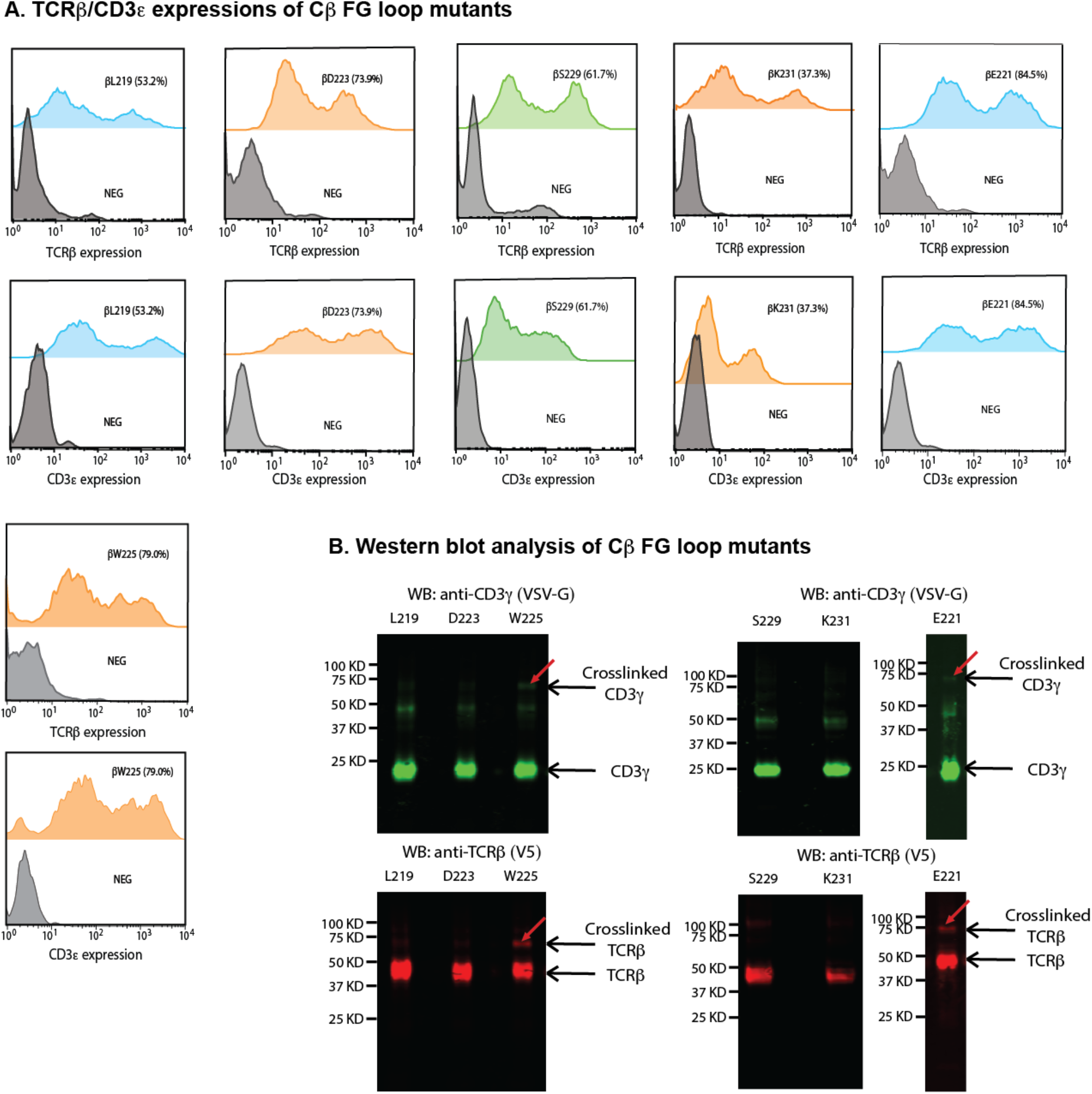
CD3γ interacts with the TCR Cβ FG loop. A) TCRβ and CD3ε expression histograms of Cβ FG loop mutants - βL219, βE221, βD223, βW225, βS229 and βK231 by flow cytometry. The percentage of cells positive for both TCRβ and CD3ε staining is indicated. B) Western blot analysis of Cβ FG loop mutants - βL219, βE221, βD223, βW225, βS229 and βK231. CD3γ crosslinked bands seen for βE221 and βW225 are apparent below 75 kDa. The blot was stained with anti-TCRβ (V5) antibody and anti-CD3γ (VSV-G). Anti-rabbit IRDye 680LT- and anti-mouse IRDye 800CW were used as secondary antibodies for detection.

**Figure 2 - figure supplement 5:**
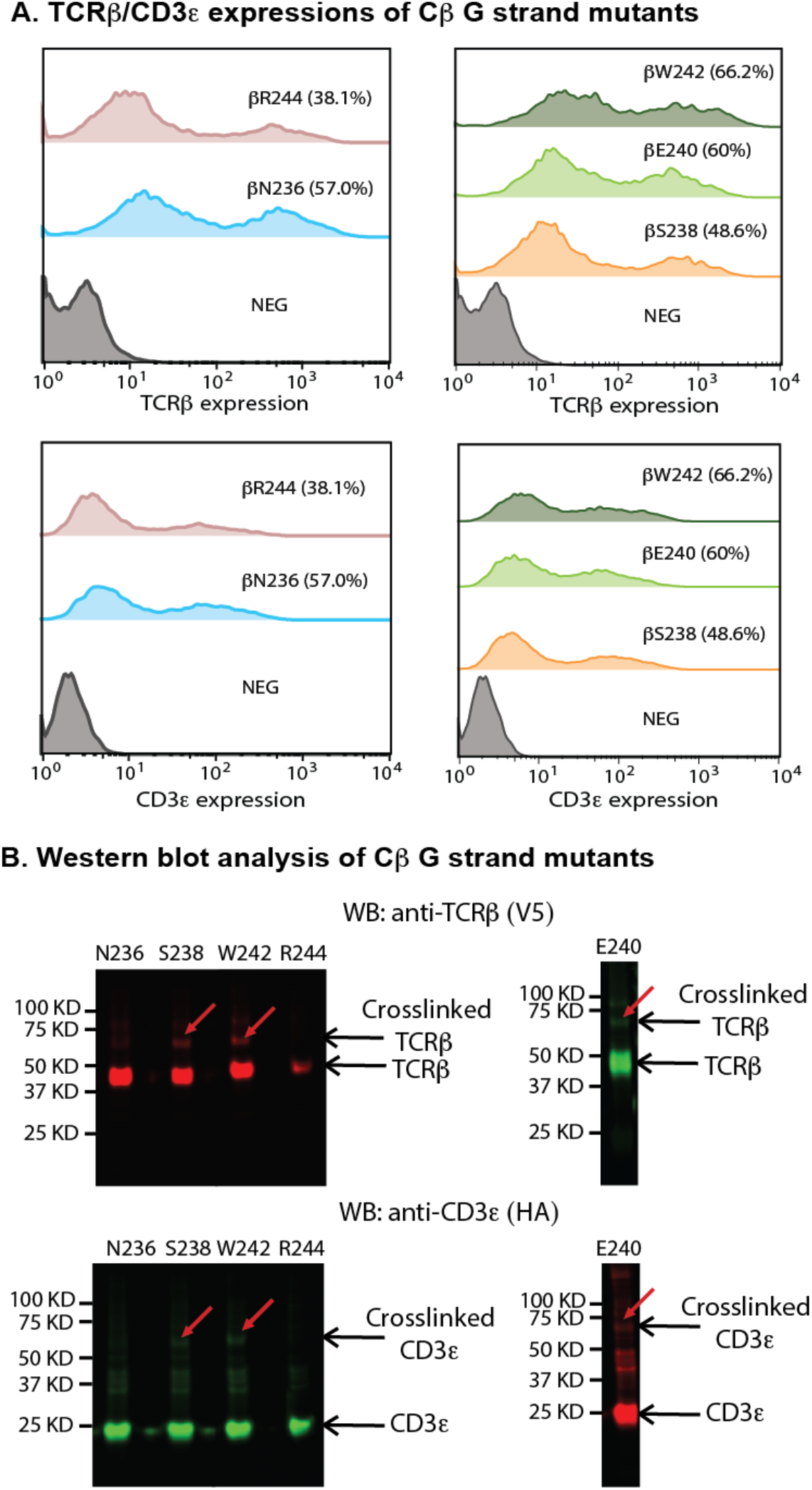
CD3ε interacts with the TCR Cβ G strand. A) TCRβ and CD3ε expression histograms of Cβ G strand mutants - β N236, βS238, βE240, βW242 and βR244 by flow cytometry. The percentage of cells positive for both TCRβ and CD3ε staining is indicated. B) Western blot analysis of Cβ G strand mutants - βN236, βS238, β E240, βW242 and βR244. CD3 ε crosslinked bands seen for βS238, β E240 and β W242 are apparent below 75 kDa. βN236, βS238, βW242 and βR244 blots were stained with rabbit anti-TCRβ(V5) antibody and mouse anti-CD3ε (HA). βE240 blot was stained with mouse anti-TCRβ(V5) antibody and rabbit anti-CD3ε (HA). Anti-rabbit IRDye 680LT- and anti-mouse IRDye 800CW were used as secondary antibodies for detection.

**Figure 2 – figure supplement 6:**
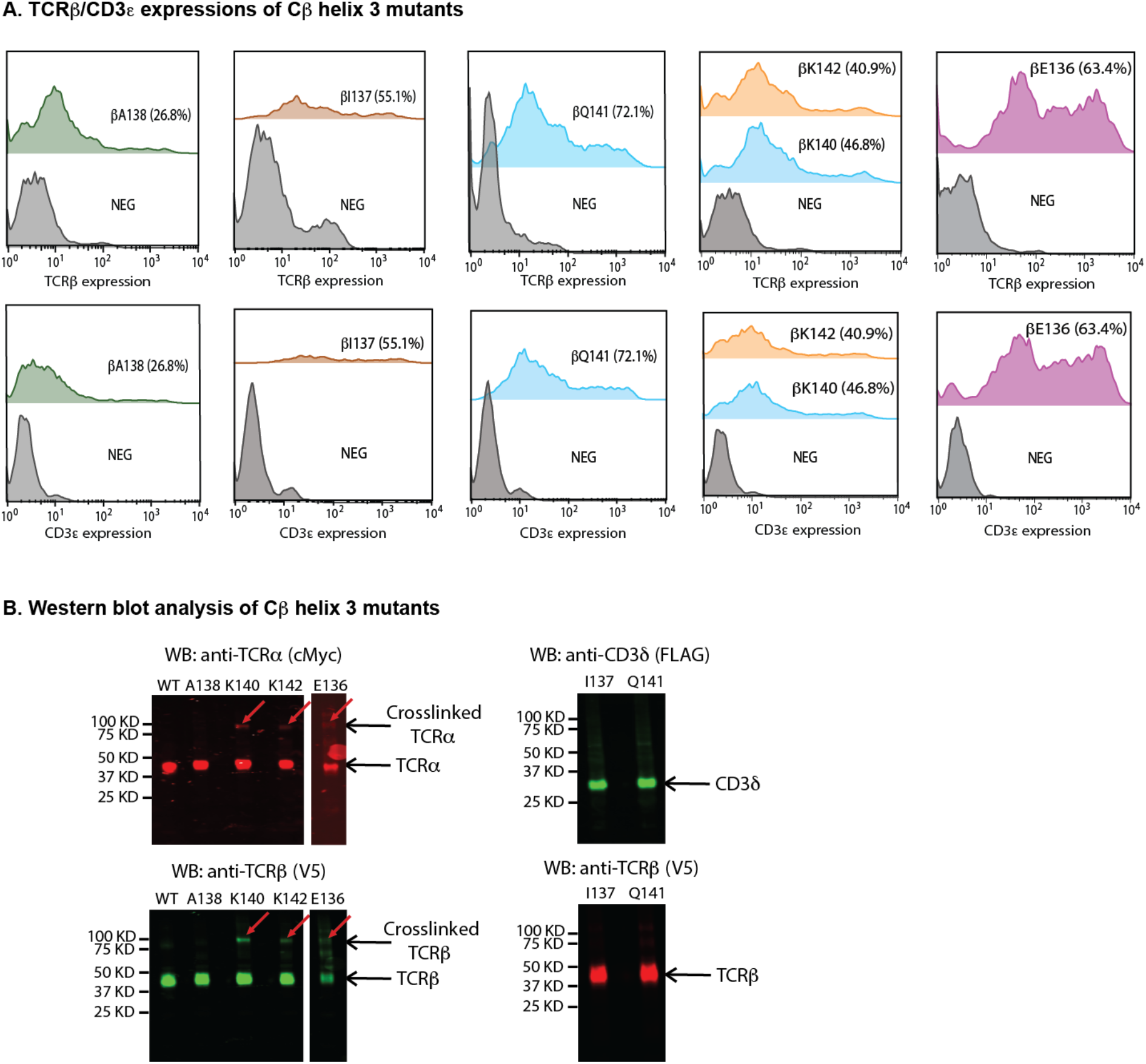
Cβ helix 3 residues interact with the TCRα subunit. A) TCRβand CD3ε expression histograms of Cβ helix 3 mutants –βE136, βI137, βA138, βK140, βQ141 and βK142 by flow cytometry. The percentage of cells positive for both TCRβ and CD3ε staining is indicated. B) Western blot analysis of Cβ helix 3 mutants - WT, βE136, βI137, βA138, βK140, βQ141 and βK142. TCRα crosslinked bands seen for βE136, βK140 and βK142 are apparent between 75 and 100 kDa. WT, βE136, βA138, βK140 and βK142 blots were stained with rabbit anti-TCRα(cMyc) antibody and mouse anti-TCRβ (V5). βI137 and βQ141 blots were stained with rabbit anti-TCRβ (V5) antibody and mouse anti-CD3δ (FLAG). Anti-rabbit IRDye 680LT- and anti-mouse IRDye 800CW were used as secondary antibodies for detection.

**Figure 2 - figure supplement 7:**
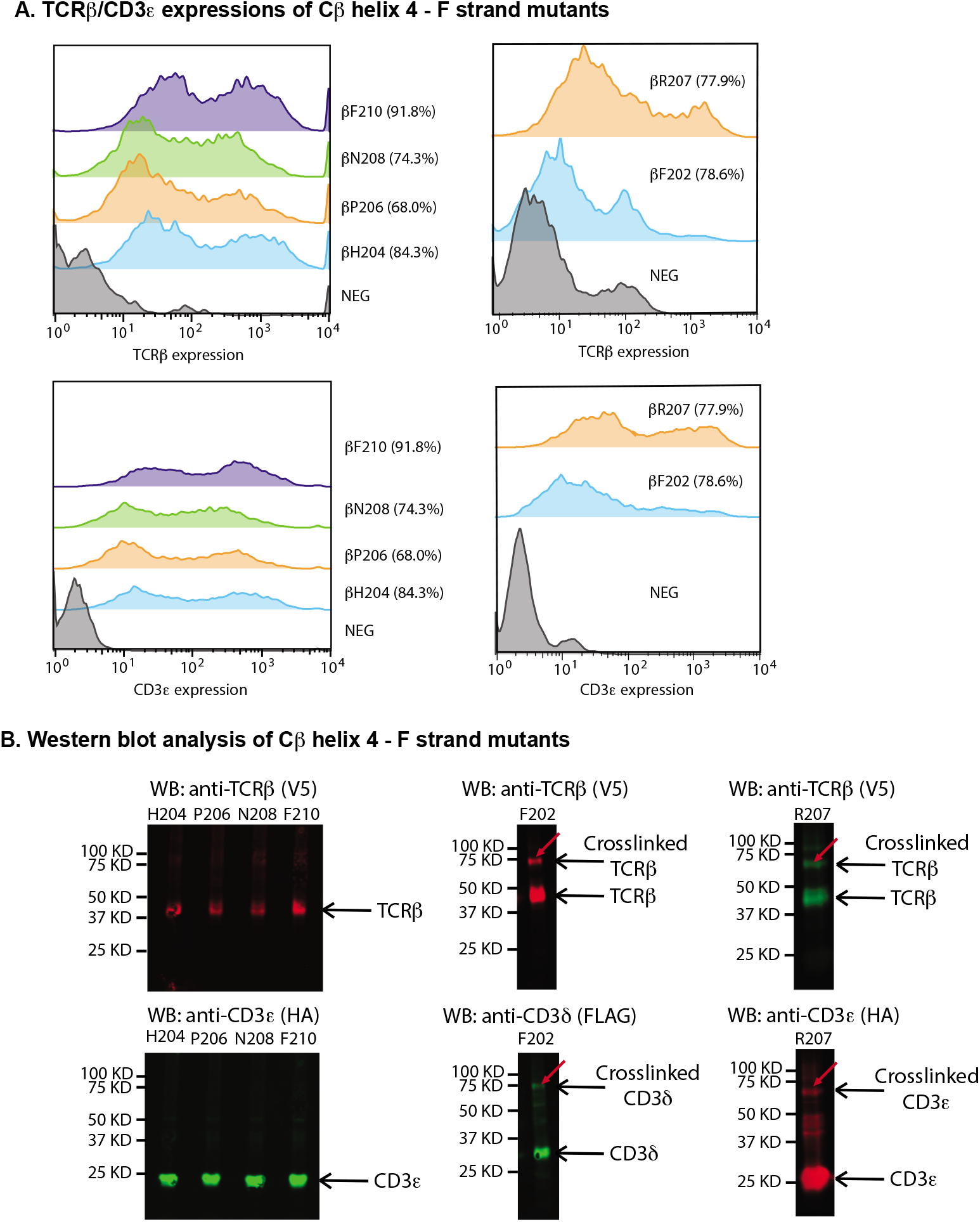
CD3δε interacts with the TCR Cβ helix 4-F strand region. A) TCRβ and CD3ε expression histograms of Cβ helix 4–F strand mutants - βF202, βH204, βP206, βR207, βN208 and βF210 by flow cytometry. The percentage of cells positive for both TCRβ and CD3ε staining is indicated. B) Western blot analysis of Cβ helix 4-F strand mutants - βF202, βH204, βP206, βR207, βN208 and βF210. CD3δand CD3εcrosslinked bands seen for βF202 and βR207, are apparent below 75 kDa. βH204, βP206, βN208 and βF210 blots were stained with rabbit anti-TCRβ (V5) antibody and mouse anti-CD3ε (HA). βF202 blot was stained with rabbit anti-TCRβ (V5) antibody and mouse anti-CD3δ (FLAG). βR207 blot was stained with mouse anti-TCRβ (V5) antibody and rabbit anti-CD3ε (HA). Anti-rabbit IRDye 680LT- and anti-mouse IRDye 800CW were used as secondary antibodies for detection.

**Figure 3 – figure supplement 1:**
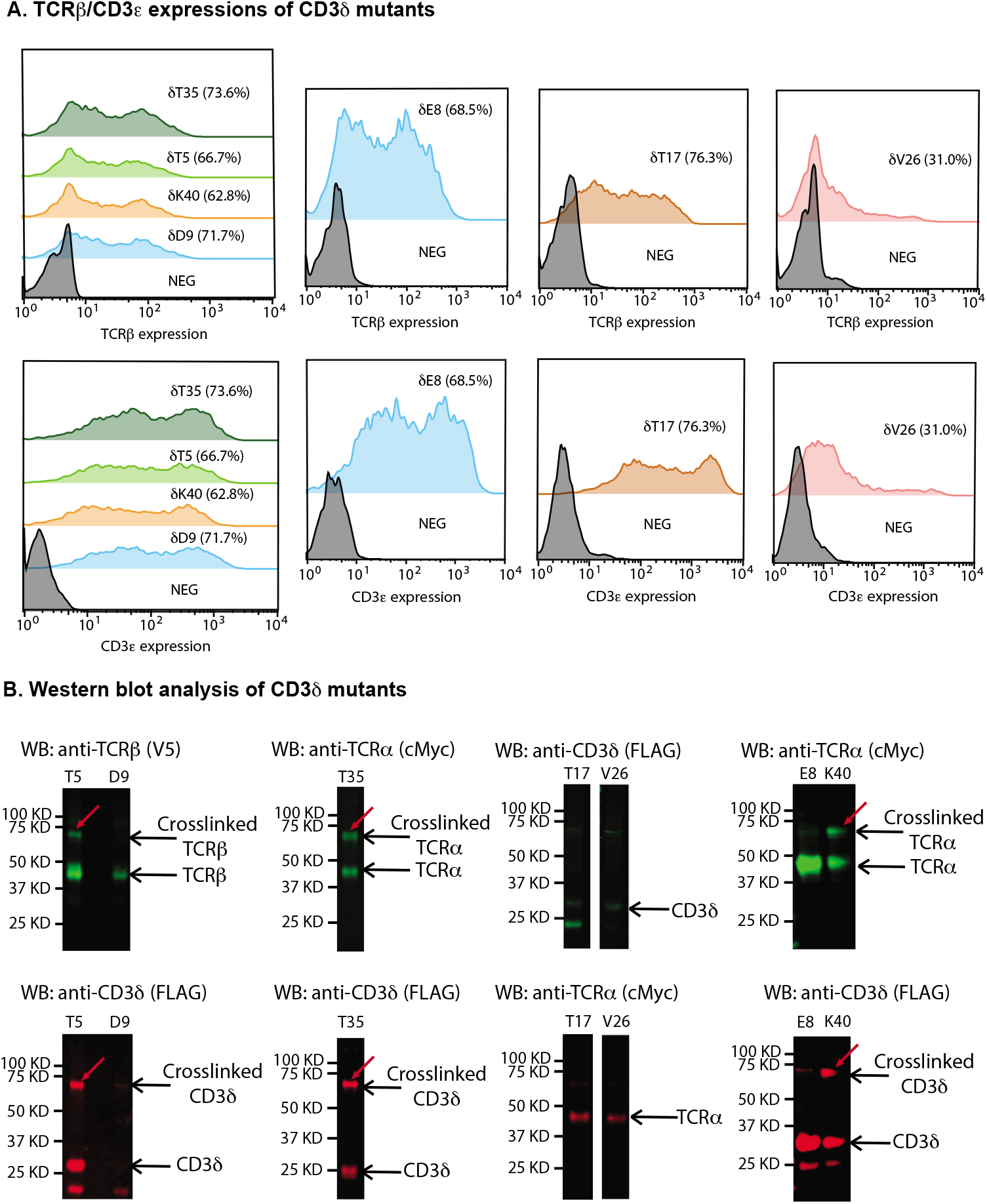
TCRα crosslinks to CD3δ E strand, CD3δ EF loop and TCRβ crosslinks to CD3δ A strand. A) TCRβ and CD3ε expression histograms of CD3δ mutants – A strand: T5; AB loop: E8, D9; BC loop: T17; CD loop: V26; E strand: T35; and EF loop: K40 by flow cytometry. The percentage of cells positive for both TCRβ and CD3ε staining is indicated. Western blot analysis revealed CD3δT5 crosslinked with TCRβ and CD3δT35 and K40 crosslinked with TCRα with crosslinking bands apparent below 75 kDa. E8, T35 and K40 were stained with mouse anti-TCRα (cMyc) antibody and rabbit anti-CD3δ (FLAG). T5 and D9 were stained with mouse anti-TCRβ (V5) and rabbit anti-CD3δ (FLAG). T17 and V26 were stained with rabbit anti-TCRα (cMyc) and mouse anti-CD3δ(FLAG). Anti-rabbit IRDye 680LT- and anti-mouse IRDye 800CW were used as secondary antibodies for detection.

**Figure 3 – figure supplement 2:**
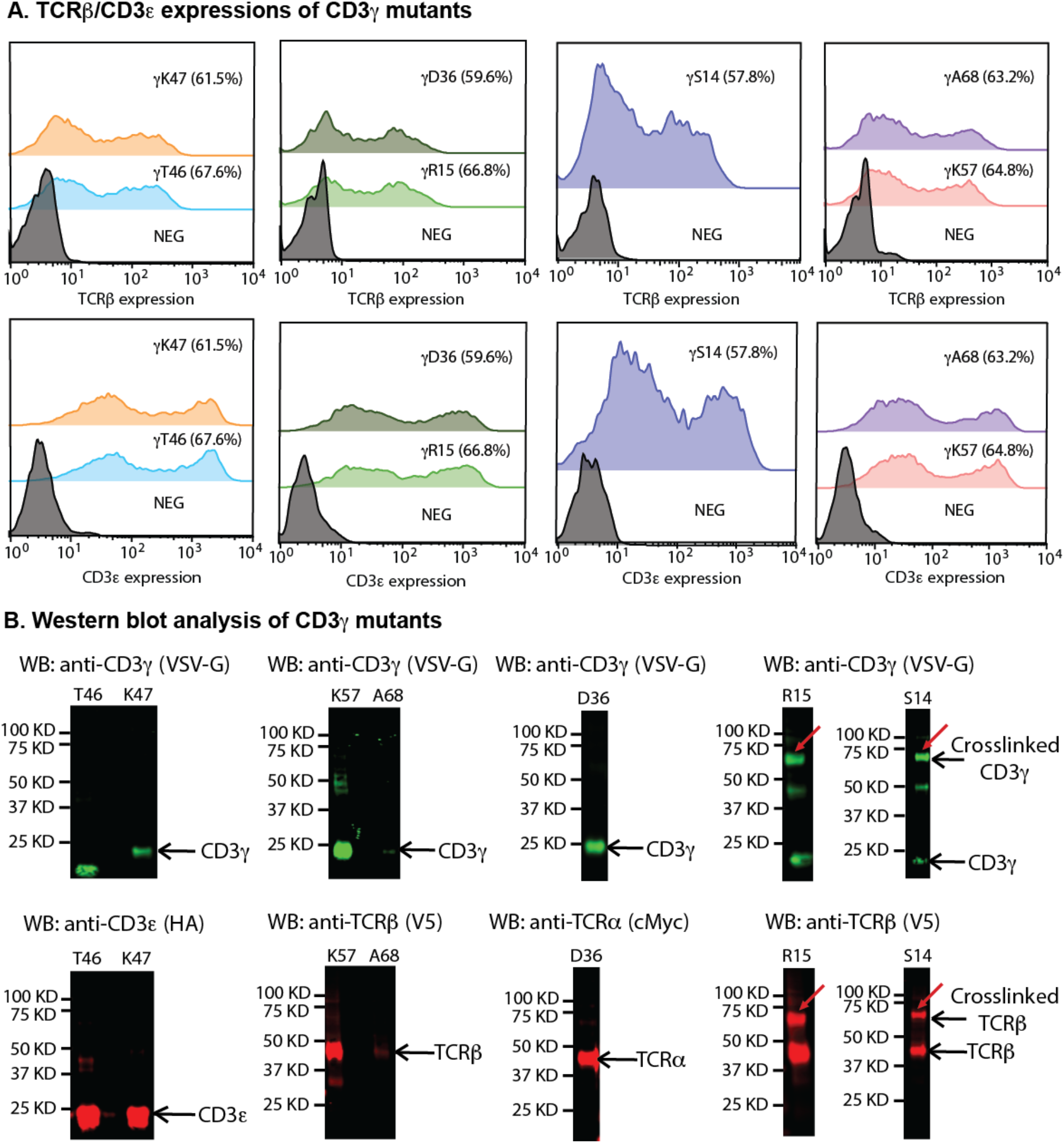
TCRβ crosslinks to CD3γ AB loop. A) TCRβ and CD3ε expression histograms of CD3δ mutants – AB loop: S14, R15; CD loop: D36; DE loop: T46, K47; EF loop: K57 and FG loop: A68 by flow cytometry. The percentage of cells positive for both TCRβ and CD3ε staining is indicated. Western blot analysis revealed CD3γ S14 and R15 crosslinked with TCRβwith crosslinking bands apparent below 75 kDa. S14, R15, K57 and A68 were stained with mouse anti-CD3γ(VSV-G) and rabbit anti-TCRβ (V5) antibody. T46 and K47 were stained with mouse anti-CD3γ (VSV-G) and rabbit anti-CD3ε (HA). D36 was stained with mouse anti-CD3γ (VSV-G) and rabbit anti-TCRα (cMyc). Anti-rabbit IRDye 680LT- and anti-mouse IRDye 800CW were used as secondary antibodies for detection.

**Figure 4 – figure supplement 1:**
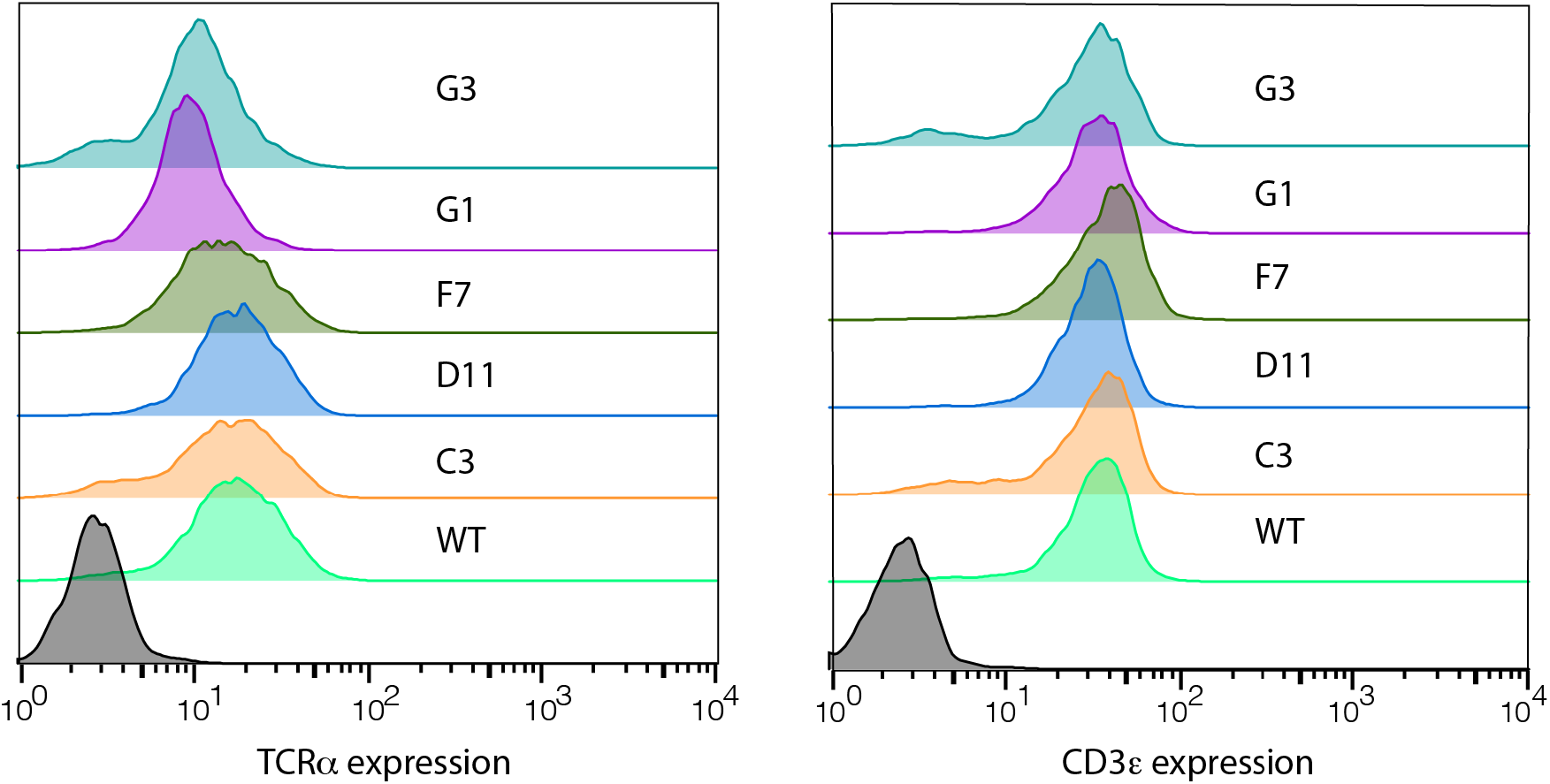
TCRα and CD3ε expression histograms of T cell hybridoma mutants tested: G3 - Cβ W242A, G1 – Cβ S238A, F7 – Cβ E221A/W225A, D11 – Cα A172G/D174A and C3 – Cβ S170A/G171A by flow cytometry.

**Figure 5 – figure supplement 1:**
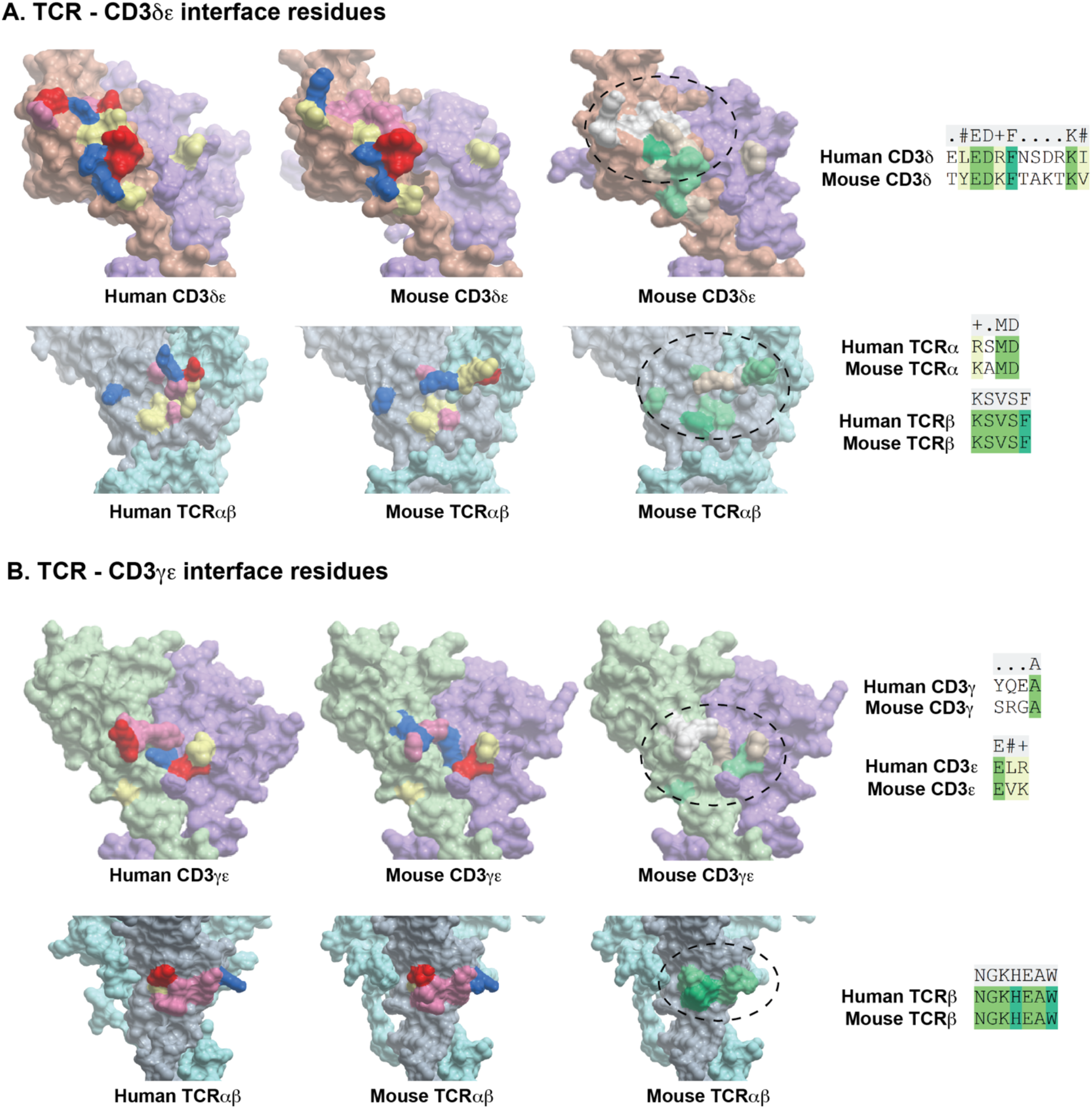
Comparison of surface charges of TCR-CD3 interface residues between human and mouse species. Positive charges are indicated in red, negative charges in blue. Green residues indicate identical residues. Overall, there is better conservation in the TCR interface between human and mouse than CD3δε and CD3γε.

**Figure 5 – figure supplement 2:**
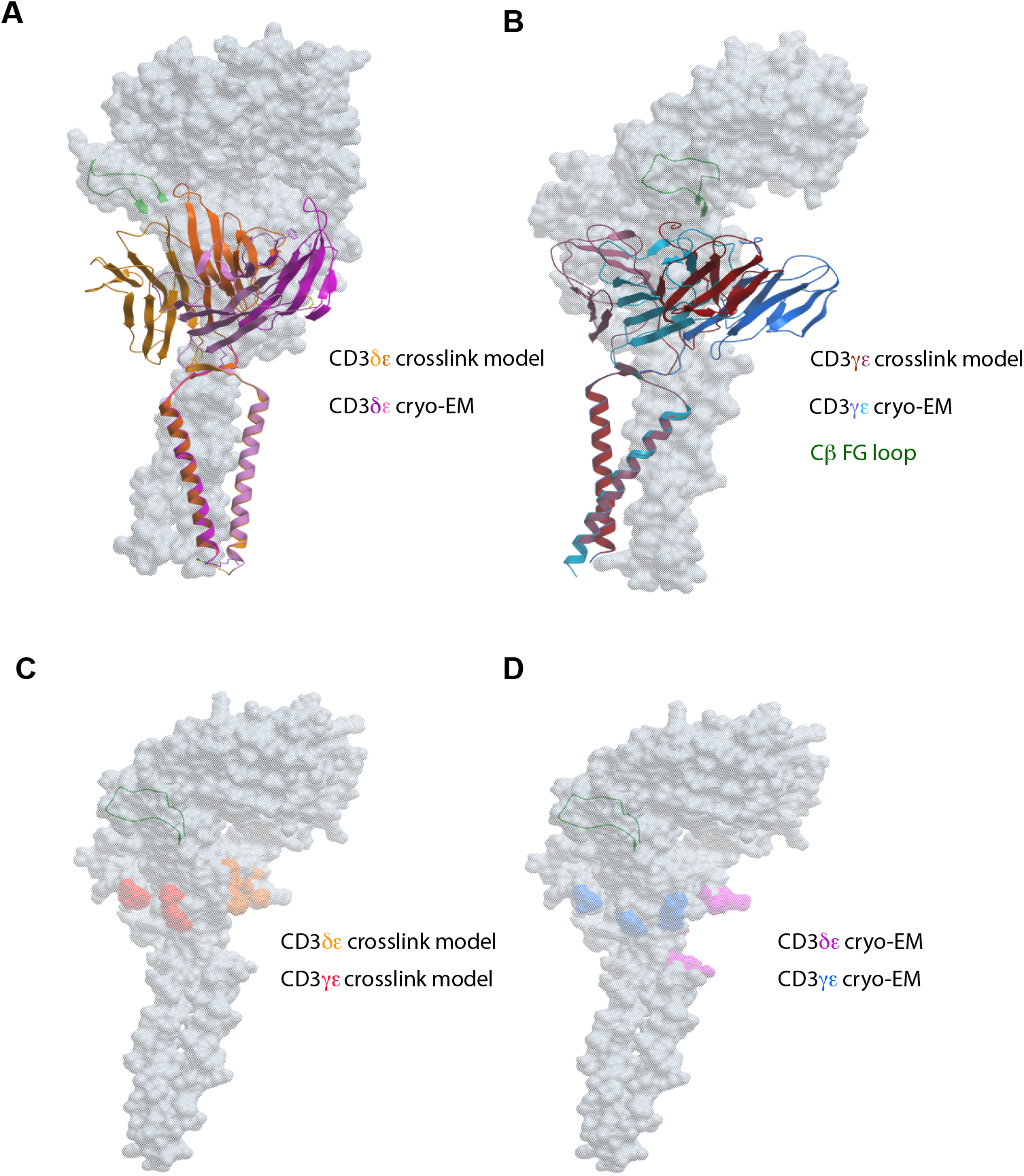
Overlay of crosslink-guided and cryoEM TCR-CD3 structures. A) Overlay of TCR-CD3δε crosslink (orange) and cryoEM (magenta) structures, B) Overlay of TCR-CD3γε crosslink (red) and cryoEM (blue) structures with TCR indicated in surface (grey) representation. C) The CD3δε (orange) and CD3γε (red) interface residues located on the TCR (grey) in the crosslink model. D) The CD3δε (magenta) and CD3γε (blue) interface residues located on the TCR (grey) in the cryo-EM structure.

## Supplementary Tables

**Figure 2 - table supplement 1:** The regions of the TCR from which specific residues were tested for crosslinking. The TCR region, sequence, speculated interacting CD3 subunits, methodology used and reference are tabled.

| Region | Sequence | Interacts with | Methodology | Reference |
| --- | --- | --- | --- | --- |
| Cβ CC' loop | <sup>164</sup>NGKEVHSG<sup>171</sup> | CD3$\gamma\epsilon$ | Mutagenesis, EM structure | (Dong et al., 2019; Kuhns and Davis, 2007) |
| Cα DE loop | <sup>170</sup>MKAMDS<sup>175</sup> | CD3$\delta\epsilon$ | Mutagenesis, EM structure | (Dong et al., 2019; Kuhns and Davis, 2007) |
| Cα AB loop | <sup>132</sup>DPRSQDS<sup>138</sup> | Conformational change upon antigen ligation | NMR, fluorescence and mutagenesis | (Beddoe et al., 2009; Rangarajan et al., 2018) |
| Cβ FG loop | <sup>219</sup>LSEEDKWPEGSPK<sup>231</sup> | CD3$\gamma\epsilon$ | NMR, antibody binding and docking | (Kim et al., 2009; Natarajan et al., 2016) |
| Cβ<br>Helix 3 | <sup>136</sup>EIANKQK<sup>142</sup> | CD3$\gamma\epsilon$ | NMR,<br>mutagenesis<br>and docking | (He et al.,<br>2015; Kim<br>et al.,<br>2010;<br>Natarajan<br>et al.,<br>2016;<br>Natarajan<br>et al.,<br>2017) |
| Cβ<br>Helix<br>4-F<br>strand | <sup>204</sup>HNPRNHFRC<sup>212</sup> | CD3$\gamma\epsilon$ | NMR, EM<br>structure,<br>mutagenesis<br>and docking | (Dong et<br>al., 2019;<br>He et al.,<br>2015; Kim<br>et al.,<br>2010;<br>Natarajan<br>et al.,<br>2016) |
| Cβ G<br>strand | <sup>232</sup>PVTQNISAEAWGRADC<sup>247</sup> | CD3ε | EM structure | (Dong et<br>al., 2019) |

**Figure 3 - table supplement 1:**
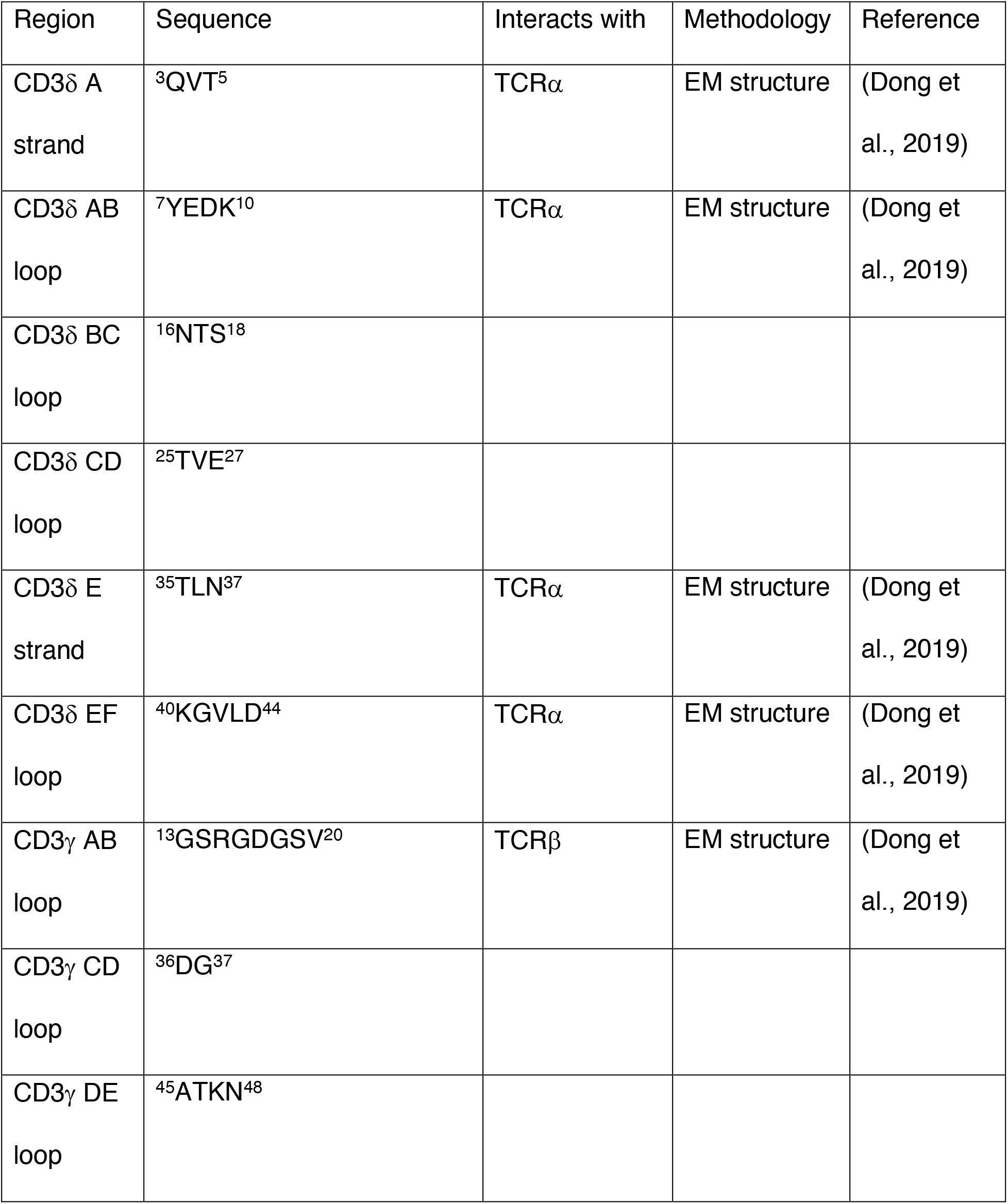
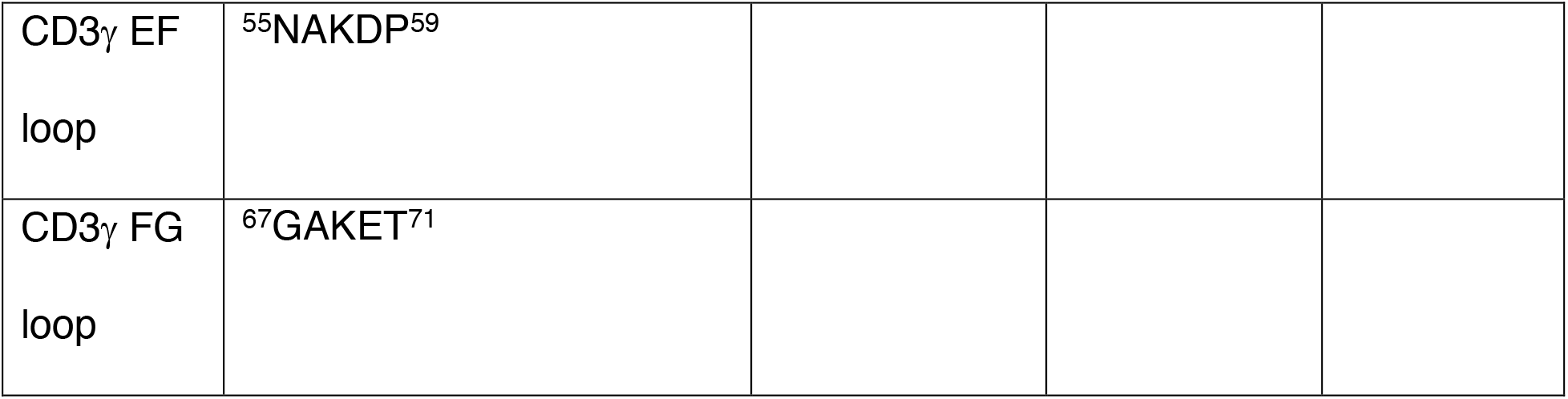
CD3 regions used for testing. The CD3 region, sequence, speculated interacting TCR subunit, methodology used and reference are tabled.

**Figure 5 - table supplement 1:** PDB structures used for the docking crosslink-guided structure.

| Subunit | Species *in vitro* | PDB | PDB Template-Species |
| --- | --- | --- | --- |
| TCR ($\alpha$ and $\beta$) V & C domains | *Mus musculus* | 3QJF and 1TCR | *Mus musculus* and *Homo sapiens* chimera; *Mus musculus* |
| TCR ($\alpha$ and $\beta$) TMs | *Mus musculus* | 6JXR | *Homo sapiens* |
| CD3$\gamma\epsilon$ | *Mus musculus* | 1JBJ | *Mus musculus* |
| CD3$\delta\epsilon$ | *Mus musculus* | 1XIW and 3R08 | *Homo sapiens* and *Mus musculus* |

**Figure 5 - table supplement 2:** Crosslinking residues to subunit distances

| CL Subunit | CL Residue | Subunit binds to | Distance (Å) |
| --- | --- | --- | --- |
| TCR $\beta$ | S238 | CD3 $\epsilon$ | 7 |
| TCR $\beta$ | E240 | CD3 $\epsilon$ | 3 |
| TCR $\beta$ | W242 | CD3 $\epsilon$ | 3 |
| TCR $\beta$ | F202 | CD3 $\delta$ | 2 |
| TCR $\beta$ | S170 | CD3 $\delta$ | 3 |
| TCR $\beta$ | G171 | CD3 $\delta$ | 3 |
| TCR $\beta$ | R207 | CD3 $\varepsilon$ | 6.5 |
| TCR $\beta$ | E221 | CD3 $\gamma$ | 6 |
| TCR $\beta$ | W225 | CD3 $\gamma$ | 7 |
| TCR $\alpha$ | A172 | CD3 $\delta$ | 5 |
| TCR $\alpha$ | D174 | CD3 $\delta$ | 7 |
| CD3 $\gamma$ | S14 | TCR $\beta$ | 5 |
| CD3 $\gamma$ | R15 | TCR $\beta$ | 8 |
| CD3 $\delta$ | T5 | TCR $\beta$ | 1 |
| CD3 $\delta$ | T35 | TCR $\alpha$ | 2 |
| CD3 $\delta$ | K40 | TCR $\alpha$ | 5 |

**Figure 2 – data supplement 1:** Mass spectrometry results indicating the proteins identified in bE221 crosslinking band. The TCR and CD3 subunits are highlighted in yellow.

**Figure 1 – source data 1:** Western blot image and files corresponding to Figure 1F.

**Figure 2 – source data 1:** Western blot images and files corresponding to Figure 2 (crosslinkers attached to the TCR regions).

**Figure 3 – source data 1:** Western blot images and files corresponding to Figure 3 (crosslinkers attached to the CD3 regions).

**Figure 4 – source data 1:** ELISA IL-2 production data for mutant T cell hybridoma and area under the curve (AUC) normalization to the wild type.

